# Protein-driven colloid-osmotic pressure controls nuclear size, organization, and function

**DOI:** 10.64898/2026.09.03.749057

**Authors:** Shin Ohsawa, Gloria Sancho-Andrés, Guido Narduzzi, Wenshuo Zhao, Dhanashree Lakhe, Marianna Estrada, Abin Biswas, Hisashi Moriizumi, Evgeny Zatulovskiy, Simone Reber, Madhav Jagannathan, Gabriel E Neurohr

## Abstract

The size of the cell nucleus is tightly controlled and changes in nuclear size correlate with altered nuclear function during development, cell differentiation and senescence^1–4^. How nuclear size is regulated^5^ and whether changes thereof are functionally relevant remains unclear. Here, we demonstrate that nuclear size is determined by the osmotic pressure exerted by proteins in yeast, human cells and frog egg extracts. The biophysical model we establish solves the long-standing question of how the nuclear-to-cytoplasmic ratio is regulated and maintained^6,7^. Furthermore, altering protein-driven osmotic balance modulates the physical properties of the nucleus, with a direct effect on chromatin organization and gene expression: Nuclear enlargement causes the dissolution of heterochromatic structures and derepression of subtelomeres and transposons, while simultaneously down-regulating highly expressed housekeeping genes. Importantly, these global transcriptional patterns closely mimic the gene expression changes that occur as yeast, human, and *drosophila* cells enlarge. Importantly, artificially forcing nuclear compression is sufficient to reverse these size-associated expression changes. Together, our findings provide a quantitative, mechanistic explanation for the coupling between the size of the nucleus and the cell, and they establish nuclear size as a modulator of chromatin organization and gene expression.

## INTRODUCTION

The size of the cell nucleus is tightly controlled. Changes in nuclear size occur during cell development, differentiation, and senescence and correlate with altered nuclear function. During development and cell differentiation, nuclear size can change substantially, and this correlates with altered transcription profiles^8,9^. In senescent human cells, increased nuclear size coincides with severe genome homeostasis defects^2,10^. Whether and how changes in nuclear size contribute to altered nuclear function remains unclear, primarily because it is not known how nuclear size is controlled.

It is clear that nuclear size is coupled to cell size. This correlation was first described in protists and developing sea-urchin embryos more than a century ago^6,7^ and has since been observed in organisms of all eukaryotic lineages^11–13^. How nuclear size is coupled to cell size is not clear. While both cell size and nuclear size are known to increase upon increased nuclear DNA content^14,15^, the coupling of nuclear size to cell size occurs independent of DNA content during early embryonic divisions and prolonged cell cycle arrests^11,12,16,17^. Genetic screens and perturbation analyses have identified nuclear transport, the cytoskeleton and the nuclear lamina as determinants of nuclear size control^18–23^. Furthermore, the nucleus responds to osmotic perturbations^21^, indicating that it is at an osmotic equilibrium with the cytoplasm. Based on these previous observations, nuclear size is considered to be governed primarily by three biophysical parameters: osmotic pressure, tension at the nuclear surface, and non-osmotic (excluded) volume^16,21,24–26^. But how this osmotic equilibrium is established and how it can explain the tight coupling of nuclear size to cell size remains unclear.

The overall osmotic pressure of the cyto- and nucleoplasm is mainly determined by small molecules such as metabolites and ions. Since small molecules with a Stokes radius of less than 2.5 nm diffuse freely across the nuclear membrane^27^, they are unlikely to contribute to an osmotic gradient between the nucleus and the cytoplasm. Instead, the osmotic pressure exerted by macromolecules such as the DNA polymer, RNA or proteins has been hypothesized to establish a fixed partitioning between nuclear- and cytoplasmic volume^16,28^. However, direct experimental evidence supporting this idea is still missing and whether the magnitude of this colloid osmotic pressure is sufficient to influence the size of organelles is unclear, as the concentration of macromolecules is two orders of magnitude smaller than that of small molecules^26^. How the osmotic pressure gradient between the nucleus and the cytoplasm is established therefore remains unknown.

Here, using massive overexpression of heterologous proteins targeted specifically to the nucleus or cytoplasm, we demonstrate that the osmotic pressure exerted by macromolecules is sufficient to alter the nuclear-to-cytoplasmic (N/C) ratio. Quantitative biophysical modeling furthermore reveals that proteins are the dominant osmolytes determining the N/C ratio in unperturbed cells. Because the ratio between cytoplasmic and nuclear proteins remains relatively stable as cells grow larger, this model explains the long-standing observation that nuclear size is coupled to cell size. Leveraging this novel system to experimentally manipulate nuclear size, we evaluated the direct consequences of nuclear scaling on nuclear function. We find that altering the colloid osmotic pressure equilibrium modulates nuclear and cytoplasmic crowding and drives a transcriptome reprogramming that strikingly mimics the gene expression changes observed during cell hypertrophy. Overall, we present a quantitative biophysical model of nuclear size determination and provide compelling evidence that nuclear size directly impacts nuclear function.

## RESULTS

### Colloid osmotic pressure generated by proteins can control nuclear-to-cytoplasmic ratio

To determine whether the osmotic pressure exerted by macromolecules is sufficient to alter the size of the nucleus, we developed an inducible, high-level over-expression system for heterologous proteins (>40 kDa) targeted to either the nucleus or cytoplasm (**Fig. 1a**). For this purpose, we expressed mScarlet-2xmCherry (3xFP) with either an SV40 nuclear localization signal (3xFP-NLS) or an HIV Rev nuclear export signal (3xFP-NES). This construct was placed under a synthetic estradiol-inducible promoter^29^ on a high copy number plasmid which can be present at up to 200 copies per cell^30^ (**Fig. 1b**). 6 hours after estradiol induction the induced proteins accounted for 8.6 ± 1.7% (3xFP-NLS) and 10.0 ± 0.6% (3xFP-NES) of total cellular protein (**Extended Data Fig. 1a, b**), and strongly partitioned to the nucleus and the cytoplasm, respectively (**Fig. 1c, d**).

**Figure 1.**
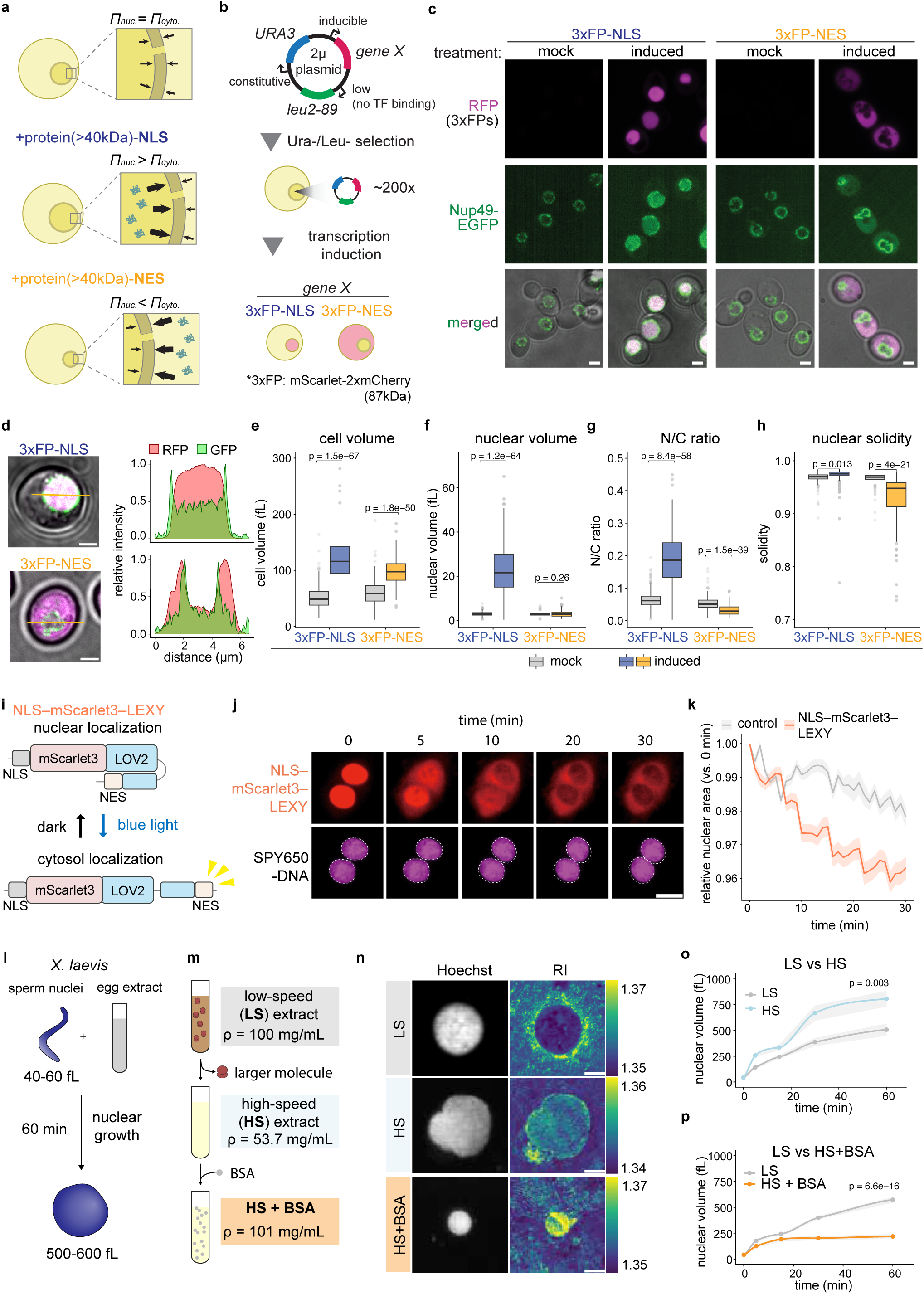
Colloid osmotic pressure generated by proteins controls the N/C ratio. **a,** Osmotic balance model between the cytosol and nucleus and predicted effect of ectopic protein expression (*Π*: osmotic pressure). **b,** Schematic of the 2µ plasmid used for estradiol-inducible expression of 3xFP-NLS or 3xFP-NES (>40 kDa). **c,** Localization of 3xFP-NLS/NES and Nup49-EGFP (nuclear envelope) after 6-h ethanol (mock) or estradiol (induced) treatment. Scale bars, 2 µm. **d,** Relative signal intensity profiles along the orange lines. Scale bars, 2 µm. **e**–**h,** Cell volume (**e**), nuclear volume (**f**), N/C ratio (**g**), and nuclear solidity (**h**) quantified from images shown in **Extended Data** Fig. 1c (mock, *n* = 894 for NLS, 602 for NES; induced, *n* = 202 for NLS, 174 for NES cells). Boxplots: center, median; box, 25th–75th percentiles; whiskers, 1.5× interquartile range (IQR); points, outliers. *P* values: two-sided unpaired *t*-test compared to mock condition. **i,** Schematic of the NLS-mScarlet3-LEXY system for blue-light-induced nuclear export. **j,** NLS-mScarlet3-LEXY localization after blue-light illumination. SPY650-DNA stained DNA. Dashed lines indicate the location of the nucleus at 0 min. Scale bar, 20 µm. **k,** Relative nuclear area after illumination. Data are mean ± s.e.m. (control, *n* = 155; OptoExport-Scarlet3, *n* = 117 cells). **l, m,** Schematics of the *X. laevis in vitro* nuclear assembly assay (**l**) and egg extract fractionation via centrifugation and subsequent supplementation with BSA (**m**). **n,** representative images of *in vitro* assembled nuclei visualized with Hoechst and dry mass density visualized using refractive index (RI) tomography. Scale bars, 5 µm. **o, p,** Nuclear volume over time after addition of sperm nuclei to LS, HS (**o**), and LS + BSA extracts (**p**). Solid lines and shaded ribbons indicate the mean and ± s.e.m., respectively (**o**, *n* = 28–37 LS, 26–39 HS nuclei; **p**, *n* = 30–201 LS, 20–30 LS + BSA nuclei). *P* values at 60 min were calculated via a two-sided unpaired *t*-test against the LS reference group.

Strikingly, we observed markedly swollen nuclei upon 3xFP-NLS induction, whereas 3xFP-NES induction resulted in an irregular, distorted nuclear morphology (**Fig. 1c**, and **Extended Data Fig. 1c**). Based on these microscopy data (**Extended Data Fig. 1c**), we segmented cells and determined the cellular and nuclear volumes. Induction of either 3xFP-NLS or 3xFP-NES led to an overall increase in cell size (**Fig. 1e**), a phenotype that was previously associated with such high levels of protein overexpression^31^. Mean nuclear volume was 7.7-fold enlarged upon 3xFP-NLS expression but remained unchanged upon expression of 3xFP-NES, despite the increased cell size (**Fig. 1f**). Consequently, the N/C ratio increased from 6.3% to 18.8% following 3xFP-NLS induction and decreased from 5.3% to 3.2% in 3xFP-NES expressing cells (**Fig. 1g**). These results demonstrate that changing the number of proteins in the nucleus or cytoplasm is sufficient to uncouple cell size from nuclear size.

In addition to altering the nuclear volume, expression of the fluorescent proteins also affected nuclear shape. Nuclear "solidity"—the ratio of the nuclear area to its convex hull area^32^—was reduced in 3xFP-NES expressing cells, consistent with the presence of nuclear invaginations (**Fig. 1h**, and **Extended Data Fig. 1d**). Conversely, 3xFP-NLS induction increased solidity (**Fig. 1h**), indicating that wrinkles in the nuclear envelope are smoothed out. Furthermore, time-lapse microscopy showed that several cells exhibited leakage of the 3xFP-NLS into the cytosol due to nuclear membrane rupture as the nuclear size increased during 3xFP-NLS induction (**Extended Data Fig. 1e-i,** and **Supplementary Video 1**). Together, these data suggest that the nuclear envelope is under little or no tension in unperturbed conditions, but expression of the 3xFP-NLS induces tension in the nuclear surface.

Massive overexpression of our constructs could have indirect effects associated with overproduction of useless proteins^31^ or by overwhelming the nuclear transport machinery. To control for such effects, we performed two control experiments. First, expressing untagged 3xFP—which partitions preferentially into the cytoplasm despite the lack of an NES (**Extended Data Fig. 2a, b**). Second, we used 1xFP-NLS which can passively diffuse through the nuclear pores and persistently burden the nuclear import system (**Extended Data Fig. 2c-e**). Overexpression of 1xFP-NLS severely impaired cell growth while induction of 3xFP-NLS, 3xFP-NES, and 3xFP caused mild and comparable growth defects (**Extended Data Fig. 2f**). These measurements suggest that the nuclear transport machinery is not overwhelmed by the 3xFP constructs and that the fitness penalty mainly stems from the production of a useless protein. Importantly, induction of 3xFP also decreased the N/C ratio (**Extended Data Fig. 2g-i**), demonstrating that biased protein partitioning alone is sufficient to perturb nuclear scaling and that this effect does not depend on altering active nuclear transport. Moreover, overwhelming the nuclear import machinery with the 1xFP-NLS construct did not increase the N/C ratio (**Extended Data Fig. 2j-l**). These controls therefore demonstrate that the N/C ratio changes upon expression of the 3xFP constructs are not driven by an overload of the active transport machinery.

The coupling of nuclear size to cell size is highly conserved and we therefore wondered whether colloid osmotic pressure also dictates nuclear size in mammalian cells. To test this hypothesis, we expressed NLS-mScarlet fused to a photo-activatable NES signal (NLS–mScarlet3–LEXY) in HeLa cells (**Fig. 1i, j**). Induction of mScarlet nuclear export led to a decrease in nuclear size by 4% over the course of 30 minutes, consistent with the idea that the osmotic pressure exerted by proteins also influences nuclear size in human cells (**Fig. 1k**).

Together, these observations show that the addition of proteins to the cytoplasm or the nucleus is sufficient to alter the shape and size of the nucleus. These experiments do not, however, distinguish whether the effect arises from the excluded volume of these proteins or from the osmotic pressure they exert. To separate the two, we varied the mean size of cytoplasmic macromolecules at constant total protein mass and assayed nuclear assembly in vitro using *Xenopus laevis* sperm nuclei and egg extract (**Fig. 1l**). Large complexes were removed by high-speed (HS) centrifugation, which lowered the total protein concentration (**Fig. 1m, n**). As previously reported ^16^, nuclei assembled in HS extract were slightly larger than those in low-speed (LS) extract, indicating that large complexes contribute to nuclear size determination, albeit modestly (**Fig. 1o, p**). We then replaced the removed mass with a small protein. Adding bovine serum albumin (BSA, 66 kDa) to the HS extract restored the original total protein mass concentration, while increasing the number of cytoplasmic colloids (HS + BSA; **Fig. 1n**, bottom). Under this condition, nuclei were smaller than in either HS or even LS extract (**Fig. 1o, p**). Nuclear size is therefore set by the number, not the mass, of cytoplasmic macromolecules, and thus by the osmotic pressure they exert across the nuclear envelope rather than by the volume they exclude.

### Quantitative modelling reveals that the nuclear-to-cytoplasmic ratio is controlled by endogenous protein osmotic pressure

The data above demonstrates that the osmotic pressure exerted by overexpressed proteins is sufficient to alter the N/C ratio, but whether endogenous proteins provide sufficient osmotic pressure to set the baseline N/C ratio remains unclear. To address this point, we generated a quantitative model that uses controlled protein induction to estimate the total number of endogenous osmotically active particles that contribute to establishing the N/C ratio (**Supplementary Note 1**). Based on the assumption that there is no tension in the nuclear envelope, the N/C Volume ratio (*r_n_*) should be proportional to the fraction of osmotically active particles that localize to the nucleus (*R_n_*)^21^ (see **Supplementary Note 1, Eq. A1.3**).

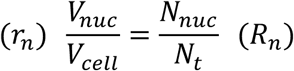

Where *N_nu_*_c_ is the number of nuclear particles and *N_t_* the total number of particles. Assuming a negligible vacuolar volume fraction (*φ_v_* = 0), we can now set the total number of particles in relation to the number of additionally introduced proteins (*n*) and with the N/C ratio *r_n_* before (*r_n,0_*) and after induction (*r_n,i_*) (**Fig. 2a**; see **Supplementary Note 1,** Eq. A1.6).

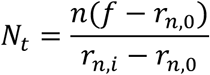

**Figure 2.**
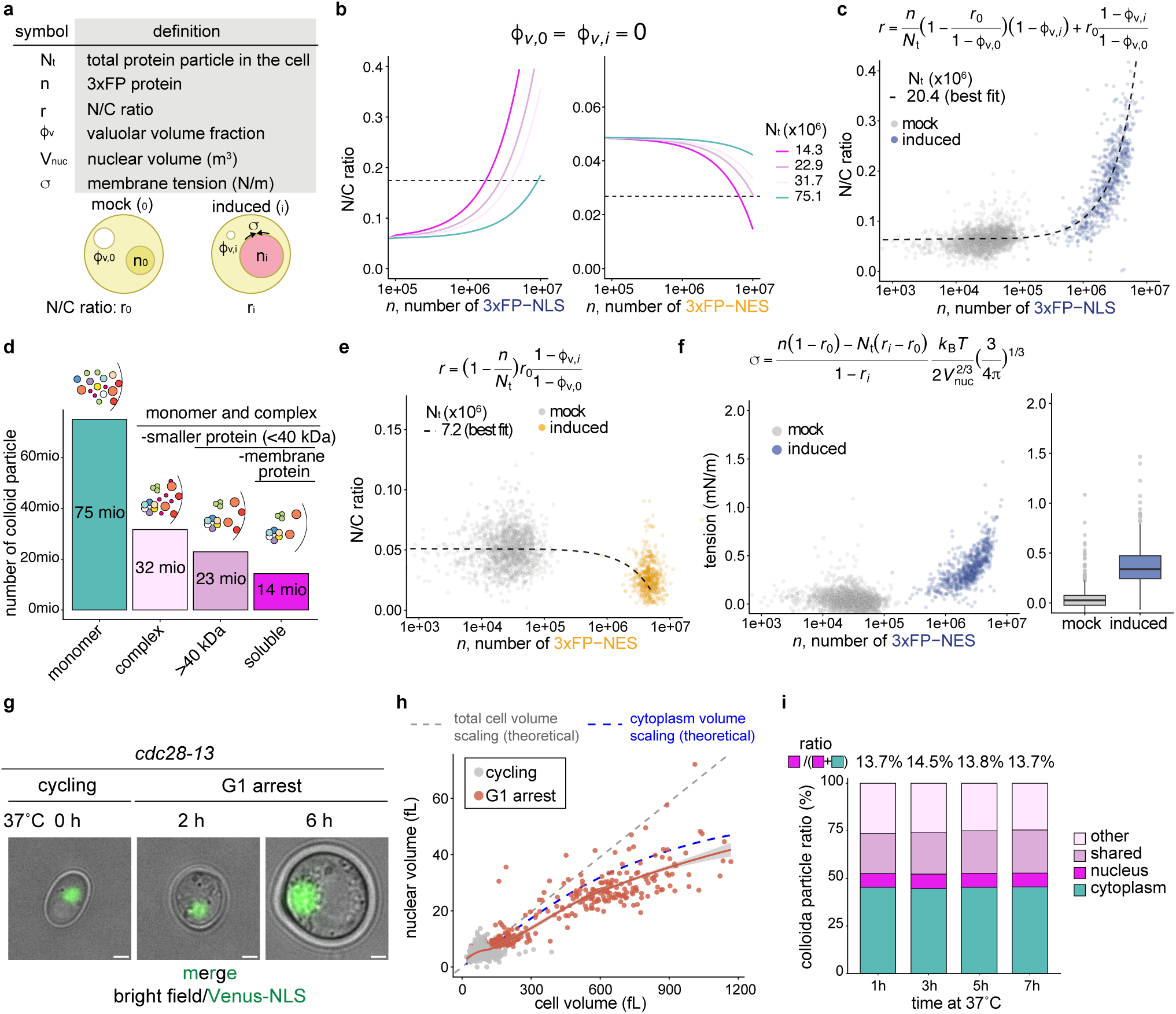
Quantitative modeling reveals the number of osmotically active particles. **a,** Definition of parameters used in the theoretical model (subscripts 0 and i denote mock and induced conditions). **b**, Prediction of how total particle number affects the response to ectopic protein expression. **c**, **e**, Number of 3xFP molecules (*n*) versus observed N/C ratios for cells expressing 3xFP-NLS (**c**) or 3xFP-NES (**e**). Sample sizes: mock (*n* = 1,744 for NLS, 1,436 for NES); induced (*n* = 685 for NLS, 524 for NES) from 3 independent replicates. Dashed lines indicate the best model fits (*N_t_* = 20.4 × 10⁶ for NLS, 7.2 × 10⁶ for NES). **d,** Estimation of protein-particle numbers based on Protein Abundance Database (PaxDb) and Complex Portal (the detail shown in *Supplementary Note 3*). Only soluble particles larger than 40 kDa are likely to contribute to influence the N/C ratio. **f**, Calculated nuclear membrane tension (*σ*, in mN/m) necessary to restrict nuclear expansion upon 3xFP-NLS induction to the observed values if we assume a constant total colloid number *N_t_* = 7.2 x 10^6^, as determined from the 3xFP-NES system. Membrane tension was thermodynamically derived using the equation shown in *Supplementary Note 4*. The dataset is the same as in **c.** Boxplots as in Fig. 1e. **g**, **h**, Representative images (**g,** scale bars, 2 µm) and correlation between cell volume and nuclear volume (**h**) of G1-arrested *cdc28-13* cells. The gray dashed line represents the theoretical perfect nuclear scaling to total cell volume using the stable N/C ratio observed in cycling cells at 30°C. The blue line shows the expected scaling between nuclear volume and cytosolic volume, factoring in the previously described disproportional increase in vacuolar fraction in very large cells^2^. The coral solid line shows the smoothed regression of the observed data. Sample sizes: *n* = 633 for cycling cells at 30°C; *n* = 280 for G1-arrested cells at 37°C. **i**, Stacked bar plots show the relative proportion of soluble protein particle (monomers and complexes ≥ 40 kDa; excluding membrane-associated proteins) localizing to the indicated compartment during the progressive cell size increase of G1-arrested *cdc28-13* cells. Proteomic data from Neurohr et al., 2019^2^. Values above the bars represent the percentage of nuclear particles relative to the total of nuclear and cytoplasmic particles.

Here, *f* represents the fraction of introduced proteins partitioning into the nucleus. Because the relative fraction of vacuolar volume slightly decreases upon expression of exogenous proteins (**Extended Data Fig. 3a**), we subtracted the vacuolar volume from the cytoplasm volume, assuming its volume is defined by small osmolytes and that it can be viewed as excluded volume in this case (**Supplementary Note 1**, Eq. A1.4). Our simulations however indicate that the observed changes in vacuolar volume have little effect on how sensitively the N/C ratio responds to expression of exogenous protein (**Extended Data Fig. 3b, c**).

Τhis formalism predicts that the N/C ratio changes faster upon induction of colloid particles if the total particle number is smaller (**Fig. 2b**). Importantly, it can be used to estimate the number of total colloid particles contributing to the N/C ratio, by fitting the model to our data. For these calculations, cell volume and total protein mass were set constant to 50 fL and 5 pg, respectively. The number of induced 3xFP molecules (*n*) was determined for each cell from the respective fluorescence intensities and the bulk 3xFP quantification (**Extended Data Fig. 1b,** and **Supplementary Note 2**). Solving equation A1.8 in Supplementary Note 1 for the induced N/C ratio (*r_n,i_*) and fitting the single cell data after 3xFP-NLS induction estimates the total number of colloids (*N_t_*) at 20.4 million (mio), corresponding to a concentration of 0.68 mM (**Fig. 2c**). This number is substantially lower than the 75 mio total proteins present in a cell estimated by proteomics^33^. But it is close to previous estimates in *Xenopus* egg extract^16^ and to the number of monomers and protein complexes larger than 40 kDa that are not associated with membranes, which we estimate to be around around 14 mio based on published proteomic data (**Fig. 2d**, and **Supplementary Note 3**).

Surprisingly, the change in N/C ratio induced by expression of 3xFP-NES suggests that the number of osmotically relevant colloids is only 7.2 mio (0.24 mM, **Supplementary Note 1**, Eq. A1.9) and thus substantially smaller than the number determined with the 3xFP-NLS construct (**Fig. 2e**). This discrepancy might be explained by the fact that the surface of the nuclear envelope could limit nuclear expansion while nuclear compression is not subject to such a restriction. To account for this possibility, we incorporated the effects of nuclear envelope tension (*σ*) into our model (see **Supplementary Note 4**, Eq. A4.4). For this purpose, we fixed the total number of colloid particles *N_t_* at 7.2 mio and estimated the nuclear envelope tension induced by nuclear expansion upon 3xFP-NLS expression. This analysis revealed that a surface tension of less than 1 mN/m (mean 0.42 mN/m) is sufficient to restrict nuclear expansion to the observed values (**Fig. 2f**). This tension is well within the range biological membranes can bear, even in the absence of a nuclear lamina^34,35^.

While nuclear envelope tension is a plausible explanation for the observed discrepancy between the measurements using the two different constructs, this difference could also have other origins, such as a reduced accuracy of measuring the volume of irregularly shaped, compressed nuclei. In either case, both measurements of endogenous colloid number are close to the estimated 14 mio colloids estimated from proteomics. These results therefore demonstrate that the relatively small osmotic pressure exerted by endogenous proteins is sufficient to determine the N/C ratio in unperturbed cells.

As mentioned above, nuclear size is tightly coupled to cell size^11,12^. This coupling can be observed during prolonged cell cycle arrests, such as those induced by the temperature-sensitive cyclin-dependent kinase allele, *cdc28-13*, during which cells continue to grow and become very large (**Fig. 2g**). In these enlarged cells, nuclear volume increases proportionally with the cytoplasmic volume (**Fig. 2h**, and **Extended data Fig. 3d**). If protein osmotic pressure is indeed the main determinant of nuclear size, the N/C ratio should be maintained as long as the partitioning of the proteome to the nucleus and cytoplasm remains constant. Analysis of our previous proteomic data in arrested *cdc28-13* cells^2^ confirmed that this is indeed the case (**Fig. 2i**). Our physical model therefore demonstrates that osmotic pressure exerted by proteins is sufficient to maintain the N/C ratio, and provides a comprehensive mechanistic explanation for the long-observed coupling of nuclear size to cell size.

### Colloid osmotic pressure determines dry mass density of the cytoplasm and the nucleus

Previous work demonstrated that the nucleus is less crowded than the cytoplasm and that this biophysical property is highly conserved^16^. Changes in crowding have important functional implications^36,37^ and we therefore wondered whether perturbing the osmotic balance between the cytoplasm and the nucleus affects their biophysical properties. Because the expressed 3xFP is smaller than the average endogenous particle (**Fig. 3a**), one would expect that its accumulation leads to a decrease in dry mass density (**Fig. 3b**). Using optical diffraction tomography, we find that the nuclear dry mass density decreases upon 3xFP-NLS induction while the dry mass density of the cytoplasm increases. Conversely, 3xFP-NES induction had the opposite effect and these reciprocal changes therefore result in a drastic shift in the N/C density ratio (**Fig. 3c, d**). Experimental perturbation of nuclear size therefore directly affects the biophysical properties of the nucleoplasm and cytoplasm, raising the interesting possibility that altering nuclear dimensions has direct consequences on nuclear function and cellular behavior.

**Figure 3.**
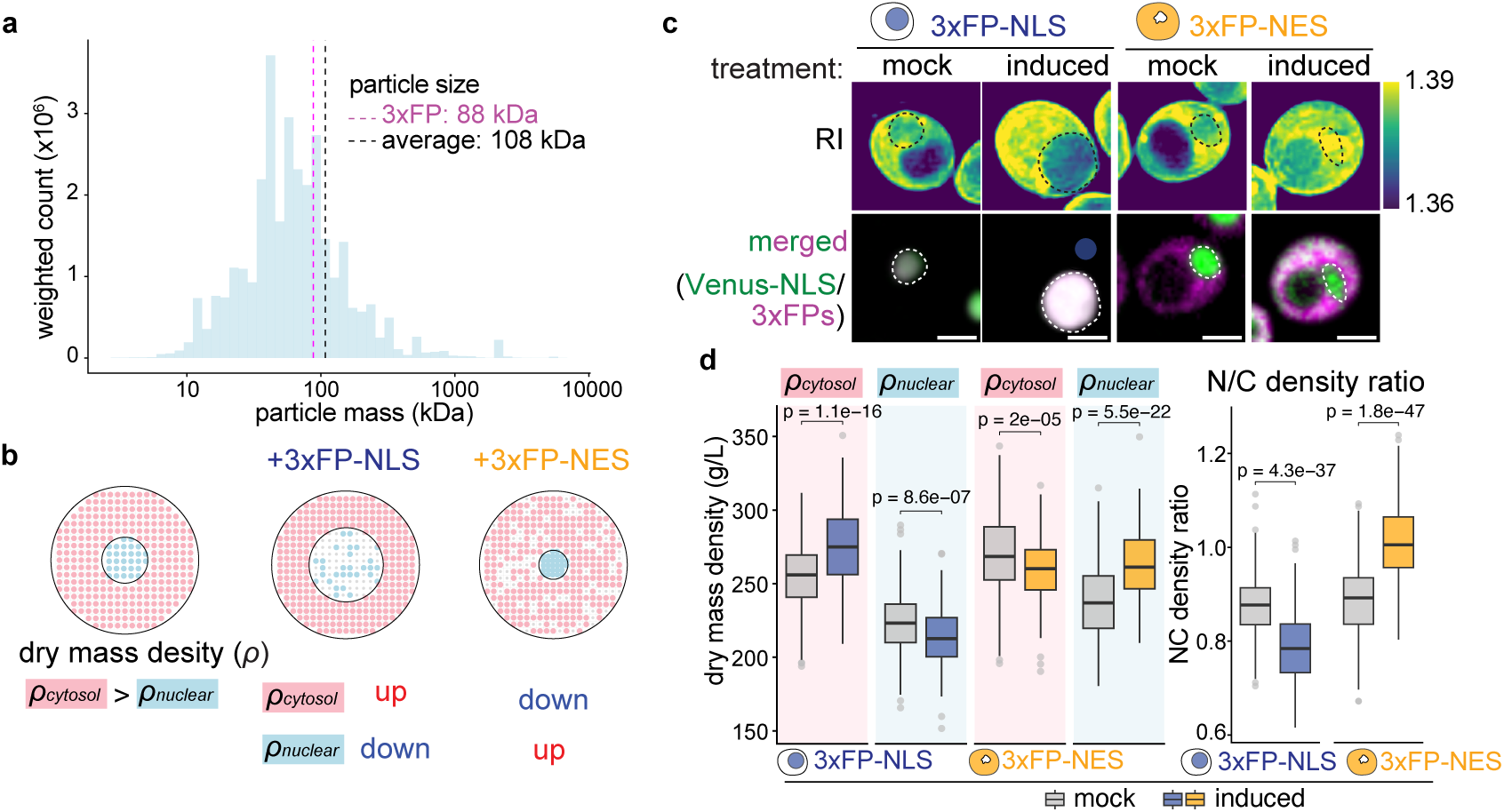
Colloid osmotic pressure controls dry mass density in the cell. **a,** Histogram of protein complex mass distribution weighted by abundance. Purple and black dashed lines indicate the mass of 3xFP and the average mass of protein particles (monomers and complexes where applicable), respectively. **b,** Schematic explaining the expected effect of altering colloid osmotic with a small particle on overall dry mass concentration. **c,** Dry mass density in yeast cells with 3xFP-NLS and 3xFP-NES induction measured using refractive index (RI) tomography. RI distribution from low (blue) to high (yellow). Dashed lines indicate the location of the nucleus. Scale bars, 2 µm. **d,** Dry mass density of the nucleus and cytoplasm were calculated from the RI values, and the nuclear-to-cytosolic (N/C) density ratio were determined. Data are from 4 replicates (mock, *n* = 240 3xFP-NLS, 240 3xFP-NES; induced, *n* = 240 3xFP-NLS, 220 3xFP-NES cells). Boxplots as in Fig. 1e. *P* values were calculated via a two-sided unpaired *t*-test.

### Experimental nuclear size changes drive global transcriptomic reprogramming

Changes in nuclear size correlate with altered gene expression during development, differentiation and cell senescence^1,19,23,38^. To what extent altered nuclear size contributes to these changes, however, remained unclear, as nuclear size remains coupled to cell size in most of these instances. Our newly developed nuclear enlargement and compression systems allowed us to directly address how altering nuclear size affects gene expression. To this end, we performed mRNA-seq analysis after induction of 3xFP-NLS, 3xFP-NES, or an empty vector. Principal component analysis (PCA) demonstrated that while mock and empty vector samples clustered together, induction of 3xFP-NLS and 3xFP-NES had distinct effects on the transcriptome (**Extended Data Fig. 4a**). These results confirm that the divergent transcriptional profiles are specifically driven by the partitioning to the cytoplasm or the nucleus, rather than by a generic response to the massive overexpression of a protein.

Plotting the differential gene expression versus the basal expression level under mock condition for each gene (MA plot) revealed substantial, bidirectional transcriptomic alterations in both samples (**Fig. 4a**). Specifically, nuclear enlargement induced a “transcriptome flattening,” characterized by the up-regulation of lowly expressed genes and the down-regulation of highly expressed genes. Conversely, nuclear compression through expression of 3xFP-NES drove a “transcriptome polarization,” wherein lowly expressed genes were further down-regulated and highly expressed genes were up-regulated. These opposing trends were statistically validated by stratifying genes into low, medium, and high expression bins (**Extended Data Fig. 4b**).

**Figure 4.**
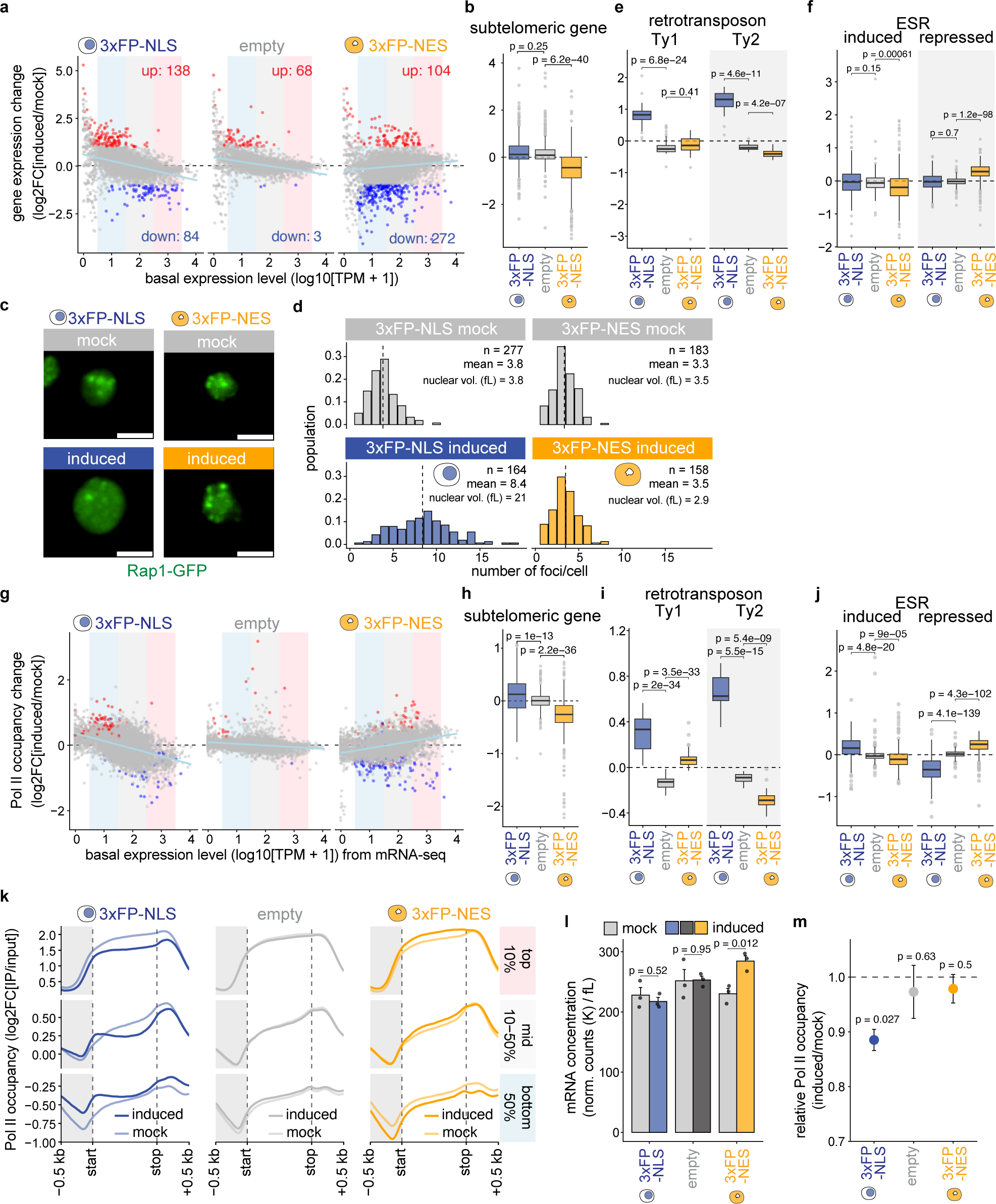
Changes in the N/C ratio shape the global transcriptome and specific genomic regions. **a**, Gene expression change (induced vs. mock) plotted against to basal gene expression level (MA plot) determined 6 h after induction of the indicated constructs. Red and blue points indicate significantly up- and down-regulated genes from three replicates (adjusted *P* < 0.05, |log2FC| > 1). Light blue lines show linear regression fits. Shaded backgrounds mark low, medium and high expression bins used for statistical analysis in Extended Data Fig. 4b. **b**, Expression changes of genes located within 30 kb of chromosome ends. Boxplots as in Fig. 1e. *P* values: two-sided paired *t*-test compared to empty plasmid control. **c**, **d**, Rap1-GFP localization (**c**, scale bar: 2 µm) and foci analysis per nucleus (**d**) after 6 h mock treatment or induction. Dashed lines indicate mean foci number. Number (*n*) of analyzed nuclei are indicated. **e**, **f**, Expression changes of Ty1 and Ty2 retrotransposons (**e**) and of environmental stress response (ESR) regulated genes (**f**) after induction of the indicated constructs for 6 h. Boxplots and statistics as in **b**. **g**, Pol II occupancy changes relative to basal RNA expression from the mRNA-seq shown in **a** for comparison. Colors indicate genes whose expression significantly changed in the mRNA-seq experiment. Light blue lines indicate linear regression. Shaded backgrounds mark is low, medium and high expression bins used for statistical analysis in Extended Data Fig. 4g. **h**, **i**, **j**, Pol II occupancy changes of genes located within 30 kb of chromosome ends (**h**), Ty1/Ty2 retrotransposons (**i**) and ESR regulated genes (**j**) 6 h after induction of the indicated constructs. *P* values: two-sided paired *t*-test compared to empty plasmid control. Boxplots as in **b**. **k**, Metagene profiles of median-centered Pol II occupancy across gene bodies, separated by basal Pol II occupancy percentiles in the mock condition. **l**, Total cellular mRNA concentration. mRNA-seq counts were normalized using *C. albicans* spike-in controls and divided by mean cell volume. Bar plots represent mean + s.e.m. *P* values: two-sided unpaired *t*-test. **m**, Global Pol II occupancy changes (induced vs. mock). Global Pol II occupancy for each sample was quantified using *C. glabrata* spike-in reads from ChIP-seq data. The dashed line indicates a ratio of 1.0. Points and error bars represent mean ± s.e.m. *P* values: two-sided one-sample *t*-test against a reference value of 1.0.

Amongst the typically repressed genes in budding yeast are the ones located close (<30 kb) to the telomeres, which were up-regulated in enlarged nuclei (3xFP-NLS) but down-regulated upon nuclear compression (3xFP-NES, **Fig. 4b**, and **Extended Data Fig. 4c**). Experimental manipulation of nuclear size therefore directly affects expression of genes in the normally silenced subtelomeric DNA. Telomere silencing is mediated by the Silent Information Regulator (SIR) complex, which clusters the 32 yeast telomeres into 3-4 superclusters at the nuclear periphery, which can be visualized with the telomere binding protein Rap1-GFP^39–42^. Upon nuclear enlargement, the number of Rap1-GFP foci significantly increased, and the foci became less intense (**Fig. 4c, d**). Conversely, nuclear compression increased foci intensity without altering foci count (**Extended Data Fig. 4d**). Together, these findings show that an increased N/C ratio triggers de-repression of subtelomeric regions which coincides with telomere de-clustering.

Furthermore, Gene Ontology (GO) analysis revealed a significant up-regulation of retrotransposons in enlarged nuclei (**Extended Data Fig. 4e**). Notably, *S. cerevisiae* lacks H3K9me, and these elements are not silenced by the SIR complex^43^. The most abundant retrotransposons, Ty1 and Ty2, were both up-regulated upon nuclear enlargement, whereas nuclear compression specifically down-regulated Ty2 (**Fig. 4e**). These findings demonstrate that retrotransposon expression—particularly Ty2—directly responds to changes in the N/C ratio, suggesting that nuclear organization is a critical determinant of transposon expression.

The massive overexpression of exogenous proteins slightly reduces cell fitness (**Extended Data Fig. 2f**). Such a slow growth phenotype has previously been associated with a stereotypic transcriptional signature, termed the environmental stress response (ESR), which is triggered by various forms of stress^44,45^. To determine to what degree the transcriptional response in our experiments is driven by activation of this response, we investigated the impact of the N/C ratio on the expression of stress responsive genes. Strikingly, while both the 3xFP-NLS and the 3xFP-NES expression induce a growth delay (**Extended Data Fig. 2f**), they affect stress regulated genes in the opposite direction (**Fig. 4f**). Together, these results show that activation of the ESR can be uncoupled from slow cell growth and they suggest that this transcriptional signature is directly linked to the N/C ratio.

ChIP-seq analysis for Rpb1, the core catalytic subunit of RNA polymerase II, confirmed that the differential gene expression measured by mRNA-seq correlates well with changes in Pol II occupancy (**Extended Data Fig. 4f**). Furthermore, analysis of Pol II occupancy relative to basal expression levels revealed the same ‘transcriptome flattening’ and ‘polarization’ signatures observed in the mRNA-seq data, and these signatures were even more pronounced (**Fig. 4a, g,** and **Extended Data Fig. 4 g**). Consistently, Pol II occupancy at subtelomeric regions, the retrotransposon Ty2, and ESR genes mirrored the transcriptomic behaviors (**Fig. 4h-j**). Upon nuclear enlargement, Pol II occupancy across the 5’-UTR, gene body, and 3’-UTR decreased in genes with initially high expression levels, whereas it increased in genes with initially low occupancy and the opposite effect was observed in the nuclear compression system (**Fig. 4k**). Taken together, these findings demonstrate that changes in the N/C ratio dictate global transcriptomic reprogramming by modulating genome-wide Pol II occupancy.

Because experimentally modulating nuclear size led to a re-distribution of RNA polymerase between highly expressed- and weakly expressed genes, we wondered how nuclear size might affect overall transcription rates and mRNA concentration. For this purpose, we used *Candida albicans* and *Candida glabrata* cells as spike-in control to normalize our mRNA-seq and ChIP-seq data, respectively. To calculate the overall mRNA concentrations, we normalized the mRNA reads to the spike-in control and divided this number by the mean cell volume. This analysis revealed that nuclear compression leads to a 24% increase in overall mRNA concentration while nuclear enlargement causes a slight decrease in mRNA concentration that does not reach statistical significance (**Fig. 4l**). When we normalize the total ChIP-seq read count to the spike-in control, we see overall less bound RNA polymerase upon nuclear enlargement (**Fig. 4m**). This decrease is stronger than the observed decrease in mRNA concentration, suggesting that the reduced transcription is compensated by increased mRNA stability as previously reported^46^. Conversely, nuclear compression did not lead to an overall increase in RNA Pol II binding, yet resulted in a 24% increase in overall mRNA concentration. This increase in mRNA concentration might be driven by a systemic stabilization of mRNA or by the redistribution of RNA Pol II from weakly expressed to highly transcribed genes.

To determine whether the transcriptional changes are influenced by an overload of the nuclear transport machinery, we analyzed 3xFP and 1xFP-NLS transcriptomes. Reducing the N/C ratio via 3xFP induction fully recapitulated the 3xFP-NES phenotype (**Extended Data Fig. 5 a–d, i, k, l**). In contrast, 1xFP-NLS, which overwhelms the nuclear transport system, yielded transcriptomic effects that were generally weaker or distinct from 3xFP-NLS (**Extended Data Fig. 5 e–h, j, m, n**). These findings confirm that the global transcriptional reprogramming is driven by N/C ratio alterations and not by an altered capacity of the nuclear transport machinery.

Together, our transcriptome analysis showed that enlarging or shrinking the nucleus is sufficient for widespread transcriptional changes driven by a re-allocation of RNA polymerase in the genome that affects both overall mRNA concentrations and relative gene expression.

### Nuclear-size-dependent transcriptomic signature is conserved during pathophysiological cellular enlargement

Senescent yeast and human cells grow very large and this size increase results in globally altered transcriptomes, severe genome homeostasis defects, and a loss of proliferative potential^2,10,46,47^. Comparison between previously published transcriptome changes of enlarged G1-arrested yeast cells^48^ (**Fig. 2g** and **Fig. 5a**) and the changes induced by 3xFP-NLS mediated nuclear expansion revealed striking similarities: repressed elements such as subtelomeric regions and Ty1/Ty2 elements were up-regulated, while highly expressed genes were down-regulated (**Fig. 5b-d**, and **Extended Data Fig. 6a–c**). Furthermore, telomeres were de-clustered in enlarged cells (**Fig. 5e, f,** and **Extended Data Fig. 6d**). Cellular enlargement therefore induces the same nuclear organization- and global transcriptome changes we observed upon artificial induction of nuclear expansion.

**Figure 5.**
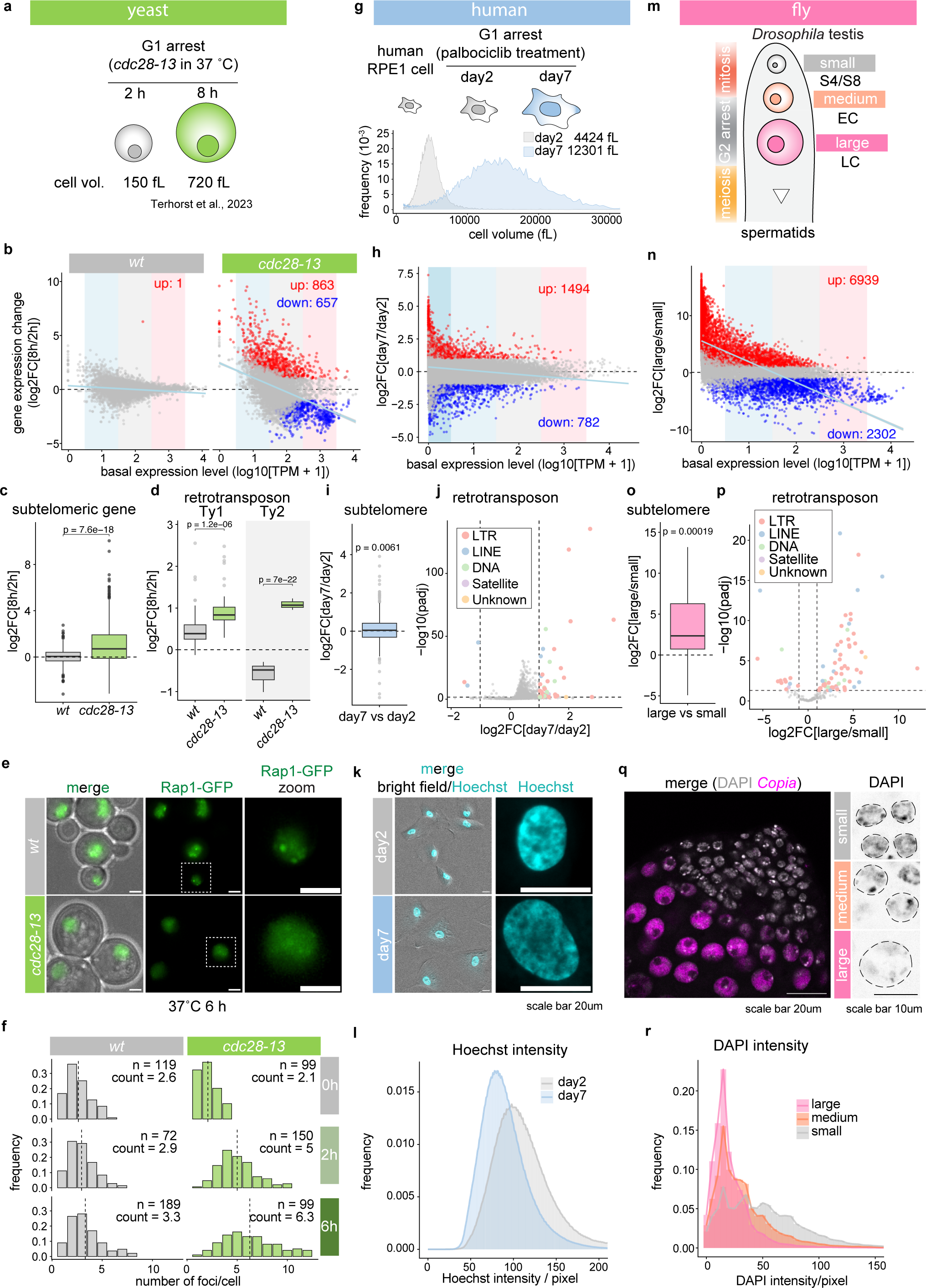
Conserved transcriptional reprogramming upon excessive cell expansion across species. **a, g, m**, Experimental models of excessive cell expansion. **a**, *cdc28-13* yeast cells during G1 arrest at 37 °C. **g**, Schematic and coulter counter volume measurements of RPE1 cells before and after a 2- and 7-day palbociclib-induced G1 arrest. **m**, Schematic of *Drosophila* spermatogenesis progressing from small germ cells (spermatogonia, S4/S8) to medium germ cell (early spermatocytes, EC) to large germ cells (late primary spermatocytes, LC) stages. **b, h, n**, Gene expression changes during cellular enlargement plotted against basal expression level (MA plot) during cell expansion. From previously published data^2^ in yeast (8 h vs. 2 h *cdc28-13* G1 arrest) (**b**). From human RPE1 (day 7 vs. day 2 palbociclib-induced G1 arrest) (**h**), and from previously published data in *Drosophila* (large (LC) vs. small (S4/S8)) (**n**). Shaded backgrounds indicate basal expression bins used for statistical analysis in Extended Data Fig. 6a, e, i. Light blue lines indicate linear regression fits. **c, i, o**, Gene expression changes of subtelomeric genes in yeast (terminal 30 kb) (**c**), human (terminal 500 kb) (**i**), and *Drosophila* (terminal 80 kb) (**o**). **d, j, p**, Transposable element (TE) expression changes. Boxplot of Ty1/Ty2 retrotransposons in yeast (**d**), and volcano plots of TEs colored by class in human (**j**) and *Drosophila* (**p**). **e, f**, Rap1-GFP localization (**e**, scale bars, 2 µm) and foci distribution quantified using spotMAX (**f**) in wild-type and *cdc28-13* cells after 6 h incubation at 37 °C. Dashed lines indicate mean foci number. Number (*n*) of analyzed nuclei are indicated. **k**, Representative images of G1-arreseted human RPE1 cells and of representative nuclei stained with Hoechst (scale bars, 20 µm). **q**, RNA FISH of the *Copia* retrotransposon and DAPI stain of a *Drosophila* testis; dashed lines outline nuclei. **l, r**, Pixel-level DNA intensity distributions (Hoechst or DAPI) for RPE1 (**l**, *n*=288 for day 2, *n*=590 for day 7) and *Drosophila* cells (**r**, *n*=25 per stage). *(General statistical note)*: For all MA and volcano plots, significant up-(red) and down-regulated (blue) genes or TEs are defined as adjusted *P* < 0.05 and |log2FC| > 1 (Wald test with Benjamini–Hochberg adjustment; thresholds indicated by dashed lines in volcano plots). Boxplot elements and statistical analyses are as in Fig. 4b, except in **i**, and **o**, where significance against 0 was determined by a one-sample *t*-test.

As in yeast, arresting human retinal pigment epithelial 1 (RPE1) cells in G1 using the Cdk4/6 inhibitor palbociclib results in excessive cell and nuclear enlargement and cell senescence (**Fig. 5g, k** and **Extended Data Fig. 6h**)^2,10^. We performed RNA sequencing and differential gene expression analysis comparing enlarged (12300 fL) to small (4424 fL) G1-arrested cells. This analysis revealed a transcriptome flattening, characterized by the up-regulation of lowly expressed genes and the down-regulation of highly expressed genes, mirroring the phenotype observed in yeast with enlarged nuclei (**Fig. 5h**, and **Extended Data Fig. 6e**). Furthermore, genes located within subtelomeric regions (**Fig. 5i**, and **Extended Data Fig. 6f**), and multiple transposable elements (TE), in particular long terminal repeats (LTRs) (**Fig. 5j**, and **Extended Data Fig. 6g**), were up-regulated in enlarged cells. DNA staining revealed that dense chromatin regions become less prominent in enlarged nuclei (**Fig. 5k, l,** and **Extended Data Fig. 6h**), raising the possibility that de-repression of weakly expressed genes is a consequence of global changes in nuclear organization. Together, these results show that transcriptomic reprogramming driven by nuclear and cell enlargement shares conserved features and closely resembles the changes observed upon experimental nuclear expansion.

### Transcriptomic reprogramming during cellular and nuclear expansion in *Drosophila* spermatogenesis

A dramatic increase in cell- and nuclear size also occurs during *Drosophila* spermatogenesis, when germ cells prepare for meiotic divisions in a prolonged meiotic G2 arrest ^49^ (**Fig. 5m**). Re-analysis of the developmental transcriptome data^50^ revealed a striking similarity between the transcriptomes of the large late spermatocytes (LC) and those in yeast and human cells with enlarged nuclei. These enlarged cells exhibited transcriptome flattening (**Fig. 5n** and **Extended Data Fig. 6i**), de-repression of genes in subtelomeric regions (**Fig. 5o** and **Extended Data Fig. 6j**), and up-regulation of numerous retrotransposons (**Fig. 5p** and **Extended Data Fig. 6k**) in comparison to smaller cells (S4/S8 spermatogonia) from the same lineage. Consistently, RNA FISH staining of the LTR retrotransposon *Copia* revealed that its expression strongly correlates with increasing nuclear size (**Fig. 5q**, and **Extended Data Fig. 6l, m**) and with the decompaction of chromatin, visualized both by DAPI staining (**Fig. 5q, r**) and by measuring the distance between two 10 kb LacO loci on chromosome 2 (**Extended Data Fig. 6n, o**). Collectively, these findings show that experimental nuclear expansion and cell enlargement result in strikingly similar transcriptome changes, suggesting that size-associated gene expression changes are largely driven by changing nuclear size.

### Nuclear compression attenuates transcriptional reprogramming in enlarged cells

To directly test whether nuclear enlargement is the driver of size-associated transcriptome changes, we expressed 3xFP-NES to slow down nuclear enlargement in arrested *cdc28-13* cells (**Fig. 6a**). 3xFP-NES expression barely affected overall cell volume, but effectively compressed the nuclei, significantly reducing both the N/C ratio and nuclear solidity (**Extended Data Fig. 7a–e**). Transcriptomic profiling of cells subjected to these conditions showed that the de-repression of Ty2 transposons was less pronounced during this G1 arrest (**Extended Data Fig. 7i**), potentially due to altered nutrient conditions during the arrest. Apart from that, nuclear compression largely reversed the transcriptomic trajectory of enlarged cells. PCA revealed that 3xFP-NES induction shifted the 4-hour transcriptional profile back toward the earlier 2-hour state (**Fig. 6b**). Accordingly, nuclear compression effectively mitigated the global transcriptomic alterations induced by G1 arrest (**Fig. 6c**, and **Extended Data Fig. 7f**), preventing the derepression of subtelomeric genes and reversing the ESR signature (**Fig. 6d, e,** and **Extended Data Fig. 7g**). Furthermore, 3xFP-NES induction successfully reduced telomere de-clustering (**Fig. 6f, g,** and **Extended Data Fig. 7h**). These results collectively establish that the majority of transcriptomic and nuclear organization changes during cellular enlargement are driven by nuclear expansion.

**Figure 6.**
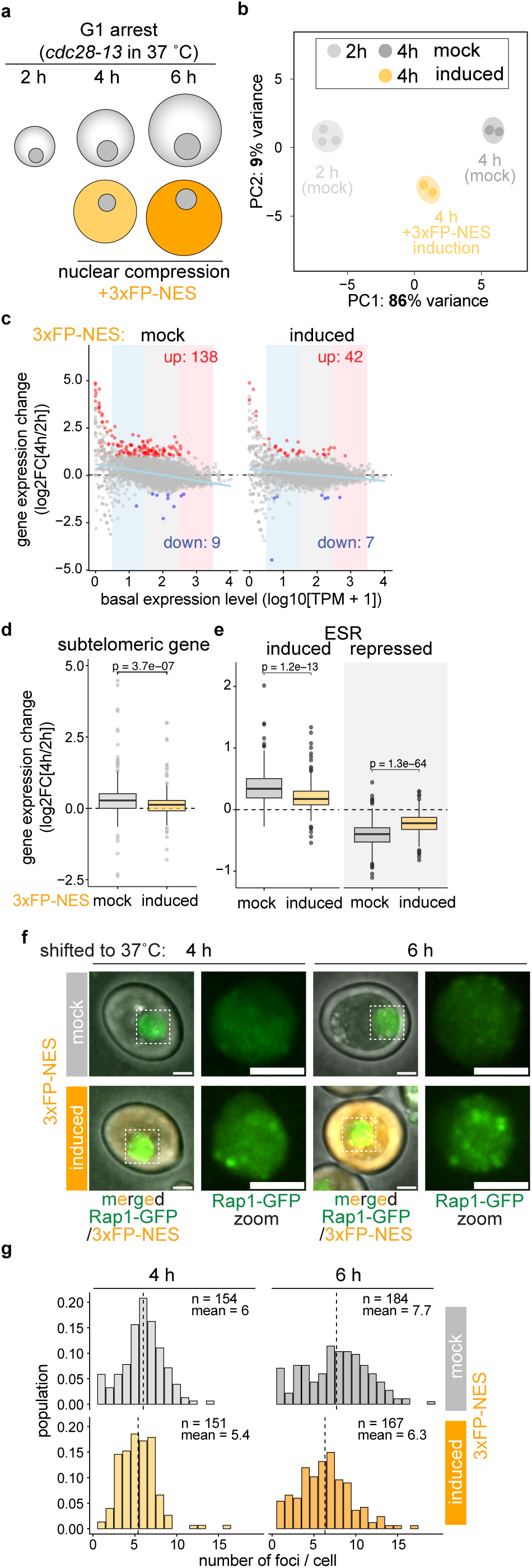
Transcriptome reprogramming in enlarged cells is mitigated by nuclear compression. **a,** Schematic of physical nuclear compression with 3xFP-NES system in G1-arrested enlarged cells. **b,** Principle component analysis (PCA) of RNA-seq data of *cdc28-13* cells arrested in G1 for the indicated times with and without induction of 3xFP-NES. **c,** Gene expression changes at 4 h relative to 2 h plotted against basal expression (MA plot). Red and blue points indicate significantly up- and down-regulated genes from three replicates (adjusted *P* < 0.05, |log2FC| > 1). Light blue lines show linear regression fits. Shaded backgrounds mark low, medium and high expression bins used for statistical analysis in Extended Data Fig. 7f. **d, e,** Gene expression change between 2 h and 4 h G1 arrest in the presence of absence of 3xFP-NES expression. Expression subtelomeric genes (terminal 30kb; **d**) and environmental stress responsive (ESR) genes (**e**) are shown. Boxplot elements and statistical analyses are as in Fig. 4b **f,** Rap1-GFP localization in *cdc28-13* with 3xFP-NES induction after 4 h and 6 h incubation at 37 °C. Scale bars, 2 µm. **g,** Distribution of Rap1 foci per cell quantified via spotMAX. Dashed lines indicate mean foci numbers. Analyzed cell numbers (*n*), and mean foci are annotated.

## DISCUSSION

Using massive protein overexpression combined with quantitative image analysis and modeling, we demonstrate that nuclear size is determined by the osmotic pressure exerted by proteins. Increasing the N/C ratio experimentally leads to transcriptional reprogramming that reflects the changes observed in enlarged cells. Compressing nuclei has the opposite effect and reverses size-associated gene expression changes. Our data explain why nuclear size is coupled to cell size, and it suggests that global transcription changes observed during cell senescence and developmental cellular enlargement are to some degree driven by changes in nuclear size.

### Osmotic pressure of proteins determines nuclear size and scaling

The coupling of nuclear size to cell size has been known since the works of Richard Hertwig^7^ and Theodor Boveri^6^. While recent work indicated that nuclear size is determined by an osmotic equilibrium between the cytoplasm and the nucleus^16,18–22^, it was not clear how this equilibrium is established as small molecules can pass freely through nuclear pores.

Our experiments demonstrate that changing the osmotic pressure exerted by proteins is sufficient to alter the N/C ratio, and they suggest that in unperturbed cells 7.2 million colloids (0.24 mM) contribute to the N/C ratio. This number is lower than the ∼14 million soluble particles estimated from proteomics. This discrepancy suggests that multiple molecules assemble into single "osmotically active particles" as suggested by previous proteomic results^16,51,52^. Regardless of the exact assembly state, this sub-millimolar concentration of colloidal macromolecules is 3 orders of magnitude lower than that of low-molecular-weight osmolytes, which was estimated to be 238 mM in yeast at by Park et al^53^. This low colloid concentration is the reason why even small forces such as the tension exerted by a biological membrane can influence the N/C ratio and explains why perturbations of the nuclear lamina affect the N/C ratio^20,54,55^.

Most importantly, our experiments demonstrate that the number of proteins in cells is sufficient to determine the N/C ratio. Intriguingly, a recent preprint by Lemière et al. comes to similar conclusions using ectopic protein overexpression in *S. pombe*^56^. This conserved colloid osmotic control system explains the mechanical basis of the universally observed nuclear size-scaling: as long as the nuclear-to-cytoplasmic protein distribution ratio is maintained, nuclear size scales linearly with cell volume during growth.

### Nuclear dimensions influence transcription

During cell differentiation and senescence, cell- and nuclear size can change dramatically and this coincides with broad changes in transcription. Using our newly developed system, we found that changing nuclear size drives the spatial redistribution of Pol II across the genome: Pol II relocalizes to weakly expressed genes during nuclear enlargement and becomes more enriched at highly expressed genes during nuclear compression.

A potential explanation for this observation is that under basal conditions, access of the transcription machinery to silent regions and low-expression genes is sterically blocked. Opening of compact chromatin in enlarged nuclei could allow binding of the transcription machinery to otherwise silenced chromatin regions. Because RNA polymerase is limiting for transcription^46^, recruitment of polymerase to normally silenced regions would titrate RNA polymerase from highly expressed genes. Indeed, accumulation of non-coding DNA in yeast nuclei is able to titrate the RNA polymerase from endogenous chromosomes, leading to a similar down-regulation of highly expressed transcripts^14^. This physical model of gene regulation could therefore explain the gene expression patterns observed upon nuclear enlargement.

But why does chromatin become more accessible in enlarged cells? Heterochromatin undergoes liquid-liquid phase separation (LLPS) and telomere clustering in budding yeast likely involves a similar process^42^. Our results show that upon cell and nuclear enlargement, dense chromatin structures become less prominent in human and *Drosophila* cells and yeast telomeres de-cluster, suggesting that chromatin condensates dissolve in enlarged nuclei. Consistent with this idea, recent work by Lemière et al show that synthetic condensates dissolve upon experimental nuclear enlargement in *S. pombe*^56^. LLPS is known to respond sensitively to changes in protein concentration and to altered biophysical properties^37,56^, which both change when nuclear size is altered. The relative contribution of these two processes to chromatin decompaction remains to be determined.

### Physiological consequences of altered N/C ratio

Nuclear size dependent changes in gene expression might have important functional consequences. In human and yeast cells, genes that change their expression disproportionally with cell size (sub- and superscaling genes) have been identified, and were proposed to serve as size sensors that allow cells to coordinate cell growth with cell division^57–59^. Similarly, activation of the environmental stress response signature has been observed in enlarged yeast cells^2,48,60^ and our data suggest that ESR activation could be a consequence of enlarged nuclear size. Nuclear size could therefore directly influence cell division, growth rate and stress resistance.

Our analysis also revealed that nuclear enlargement induces transcriptome changes that have been associated with cell senescence in yeast and human cells. De-repression of retrotransposons has been suggested to contribute to genome instability in senescent cells and to drive organismal aging and various age-related pathologies^61–63^. Similarly, loss of telomere protection has been causally linked to genome instability in senescent cells^64^. Nuclear enlargement therefore may play a key role in driving genome instability during senescence.

Finally, changes in nuclear size could facilitate developmental changes in gene expression. During spermatogenesis in *Drosophila* for example, the normally inactive Y chromosome experiences significant transcriptional induction while certain transposable elements are also expressed^65–69^. The massive cell size increase that occurs during spermatogenesis in *Drosophila* could mechanically facilitate this process, potentially through de-repression of transcriptionally silent loci^66^. However, the relative contribution of increasing nuclear size and altered activity of other TE silencing pathways, e.g. the piRNA pathway^70,71^, remains to be determined. Supporting the idea that nuclear size influences cell fate, the accompanying paper by Moriizumi et al shows that the N/C ratio is a predictor for the expression of developmental genes and differentiation trajectory in mouse embryonic stem cells^55^. Changes in nuclear size therefore might play a key role in determining gene expression, cell function and cell identity during adaptation to new environments, differentiation and senescence. How global size dependent processes and classical transcription regulation by transcription factors influence each other remains to be determined.

Taken together, our results provide a mechanistic explanation for the long-observed coupling between nuclear size and cell size. Furthermore, we show that changes in nuclear size directly affect the physical properties and gene expression, with important implications for cell function and cell identity. Our work therefore highlights the profound impact of physical regulation and weak macromolecular forces on nuclear organization and global transcriptional control.

## Acknowledgements

We thank M. Peter for sharing reagents and laboratory infrastructure, and A. Taddei for kindly providing the Rap1-GFP strain. We acknowledge the Scientific Center for Optical and Electron Microscopy (ScopeM) and the Functional Genomics Center Zurich (FGCZ) for their support with microscopy and ChIP-seq, respectively. We are grateful to Michele Marass for providing editorial advice during the publication process. This work was supported by SNSF grants (PCEFP3_187003, 320030-232190) awarded to G.E.N., and (310030_219360, 320030_228043) to M.J. S.R. acknowledges funding from the DFG (528483508 – FIP 12) and the MPG. G.N. is supported by a PhD scholarship from the Boehringer Ingelheim Fonds. E.Z. is supported by a UK Medical Research Council Career Development Award (MR/X020290/1).

## Author contribution

Conceptualization: S.O., G.S., and G.Ne.; Funding Acquisition: E.Z., S.R., M.J., and G.Ne.; Project Administration: S.O., G.S., and G.Ne.; Supervision: G.Ne.; Investigation: S.O., G.S., W.Z., D.L., M.E., A.B., and H.M.; Methodology: S.O., G.S., M.E., A.B., H.M., and G.Ne.; Formal Analysis: S.O., G.S., G.Na., W.Z., and D.L.; Software: S.O., G.Na., and W.Z.; Validation: S.O.; Data Curation: S.O.; Visualization: S.O.; Writing – Original Draft: S.O. and G.Ne.; Writing – Review & Editing: all authors.

## Data and code availability

Sequencing data from mRNA-seq and ChIP-seq have been deposited at GEO (accession number GSE339383) and are publicly available as of the date of publication. This paper also analyzes existing, publicly available data. Their accession numbers are listed in the key resources table.

All original code has been deposited on GitHub and is publicly available at https://github.com/shinohsawa/Ohsawa_et_al_2026

## Figure legends

**Extended Data Figure 1.**
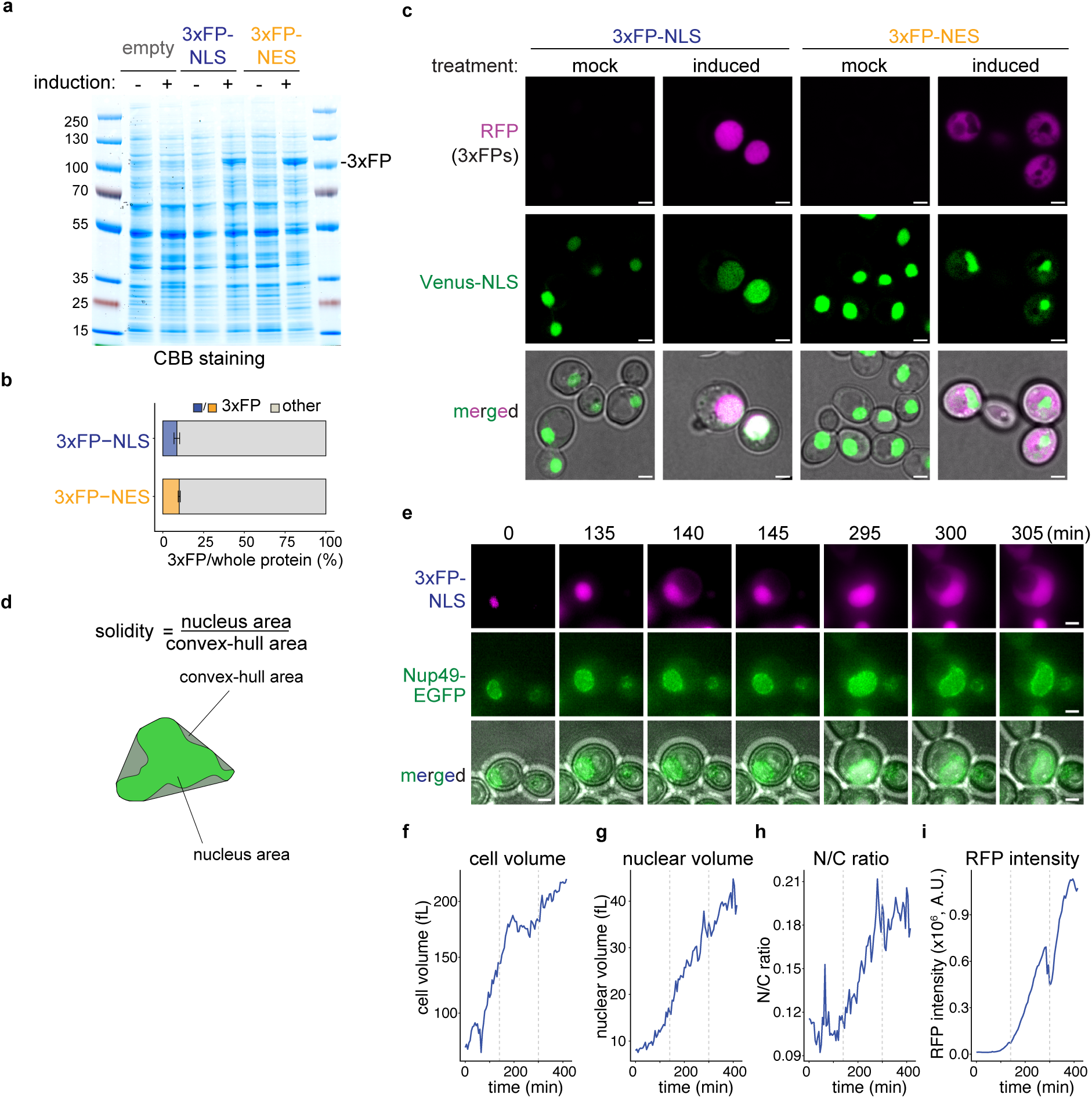
Colloid osmotic pressure generated by proteins controls N/C ratio, related to Figure 1. **a**, Equal volumes of lysate after 6-hour induction were run on SDS-PAGE followed by coomassie brilliant blue (CBB) staining. Cells containing 3xFP-NLS, 3xFP-NES, and empty vector lacking 3xFPs. **b**, Quantification of (**a**) to determine the proportion of 3xFP-NLS and 3xFP-NES to whole protein from three biological replicates. Bar plot represents mean ± s.d. **c**, Localization of 3xFP-NLS/NES and Venus-NLS (nuclei) after 6-h ethanol (mock) or estradiol (induced) treatment. Scale bars, 2 µm. **d**, Schematic defining nuclear solidity. **e**-**i**, Time-lapse images after 3xFP-NLS induction showing leakage of the fluorescent protien into the cytoplasm at 140 minutes and at 300 minutes, indicating nuclear rupture events. Nuclei were visualized with Nup49-EGFP. Scale bar, 2 µm. Cell volume (**f**), nuclear volume (**g**), N/C ratio (**h**), and magenta fluorescence intensity normalized to cell volume (**i**) in the 3xFP-NLS induced cell. Gray dashed lines indicate the time points of observed nuclear envelop ruptures (**e**; see also **Supplementary Video 1**).

**Extended Data Figure 2.**
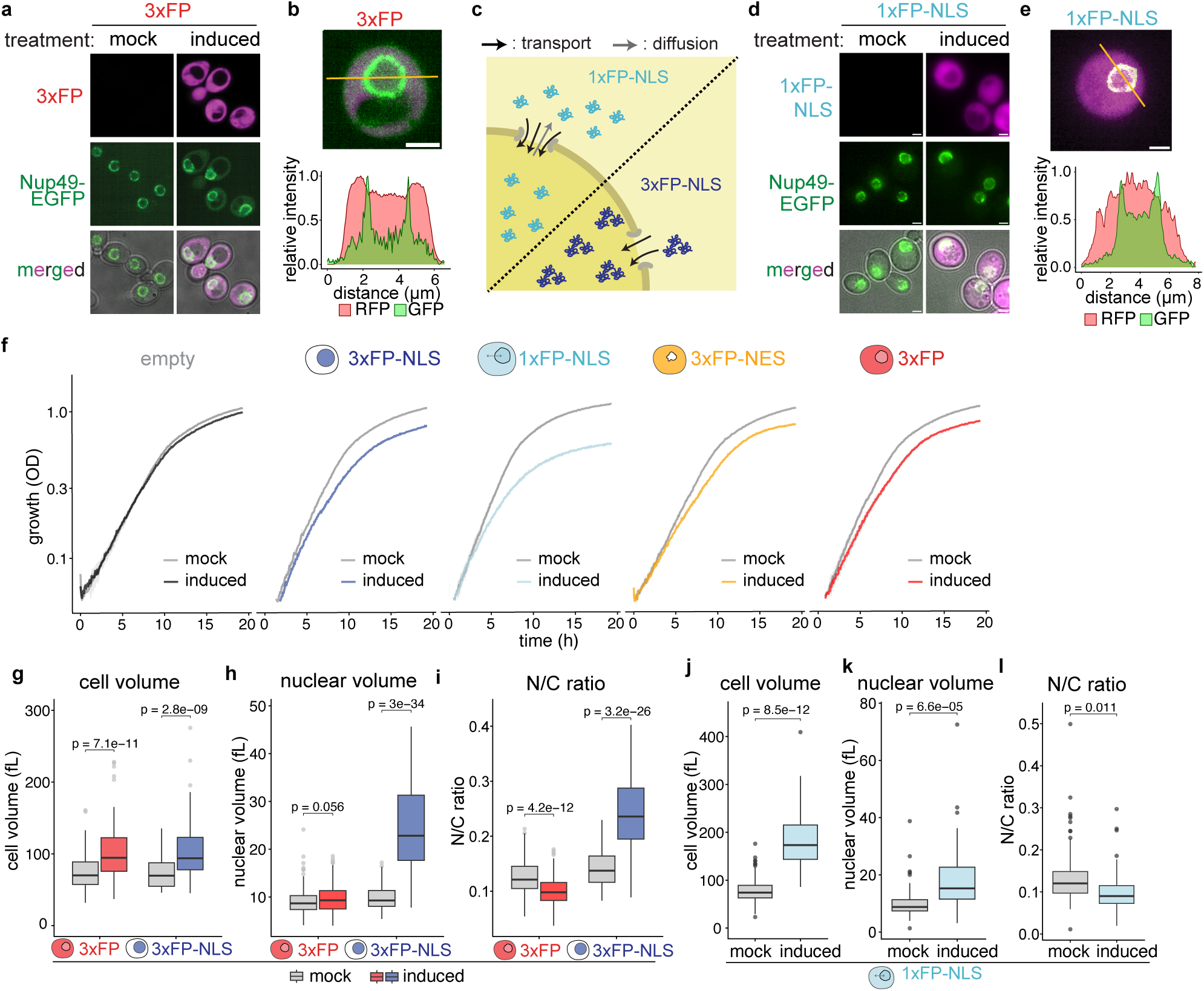
Perturbations of the N/C ratio are independent of active nuclear transport overload. **a**, Images of cells expressing of mScarlet-2xmCherry (3xFP) for 6 h compared to mock (EtOH) treated cells. **b,** Signal intensity profiles in a cell expressing 3xFP along the orange lines normalized to the maximal pixel intensity along the profile. **c**, Schematic illustrating how passive diffusion through nuclear pores (grey arrows) of small proteins with a nuclear localization signal increases the burden on the nuclear transport machinery. **d**, Images of cells expressing mCherry-NLS (1xFP-NLS) for 6 h compared to mock (EtOH) treated cells. **e**, Signal intensity profiles in a cell expressing 1xFP-NLS along the orange lines normalized to the maximal pixel intensity along the profile. **f**, Growth curve of cells containing the indicated inducible constructs in the presence of estradiol (induced) or ethanol (mock). Solid lines and shaded ribbons indicate the mean and ± s.d., respectively, across three independent replicates. **g**–**i,** Cell volume (**g**), nuclear volume (**h**), and N/C ratio (**i**) quantified from images in (**a**) (mock, *n* = 162 for 3xFP, 44 for 3xFP-NLS; induced, *n* = 134 for 3xFP, 130 for 3xFP-NLS cells, one representative dataset of three biological replicates is shown). **j**-**l**, Cell volume (**j**), nuclear volume (**k**), and N/C ratio (**l**) quantified from images in (**d**) (mock, *n* = 147 for; induced, *n* = 38 for 1xFP-NLS cells, one representative dataset of two biological replicates is shown). **a, b, d, e,** Scale bars, 2 µm. **g-l**, nuclei is segmented with Nup49-EGFP.

**Extended Data Figure 3.**
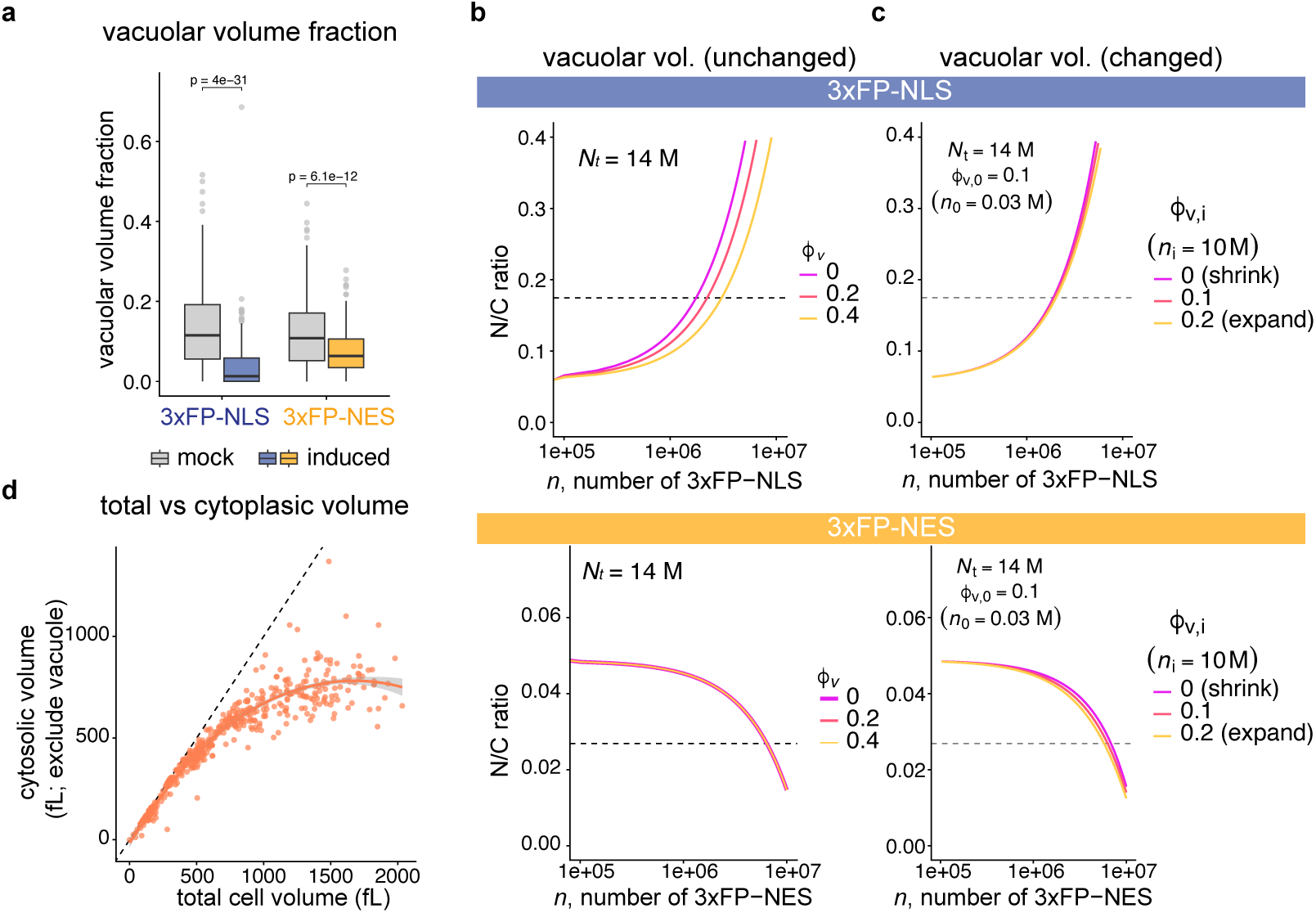
Modeling of the N/C ratio as a function of 3xFP levels and vacuolar fraction. **a**, Vacuolar volume fraction after induction of the indicated constructs using was determined using refractive index tomography (see Fig. 3c). Data are pooled from three independent experiments (mock, *n* = 242 for 3xFP-NLS, 238 for 3xFP-NES; induced, *n* = 250 for 3xFP-NLS, 266 for 3xFP-NES cells). *P* values denote comparisons between mock and induced treatments for each strain, calculated via a two-sided unpaired Student’s *t*-test without adjustment. **b**, **c**, Theoretical predictions of effects of vacuolar volume fraction on how the N/C ratio changes upon addition of proteins to the cytoplasm or the nucleus. (**b**) shows the prediction for a constant vacuolar volume fraction and (**c**) shows a scenario where the volume fraction changes from 0.1 to the indicated values. Boxplot elements and statistical analyses are as in Fig. 1e. **d**, Scatter plot plotting the cytosolic volume (exclude vacuole) against the total cell volume. The cytosolic volume was estimated by quantifying the intracellular regions containing the freely diffusing cytosolic marker Pgk1. Each point represents an individual cell (*n* = 518 cells). The solid coral line indicates the smoothed conditional mean trend, and the dashed gray line represents the identity line (*y* = *x*). Raw image data were adapted from Neurohr et al., 2019^2^.

**Extended Data Figure 4.**
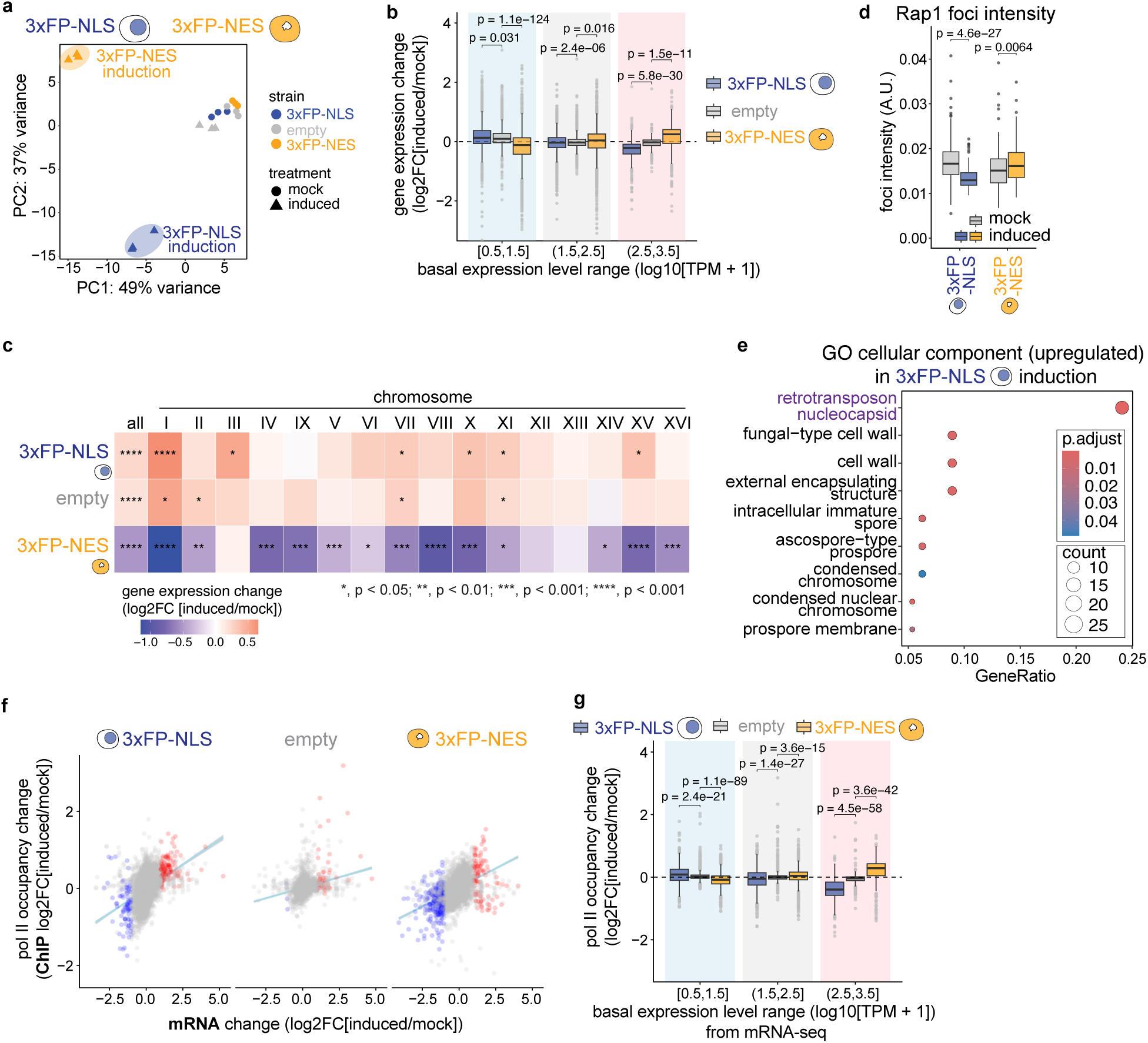
Changes in the N/C ratio shape the global transcriptome and specific genomic regions, related to Figure 3. **a**, Principle component analysis (PCA) of RNA-seq data of strains containing the indicated construct after mock (EtOH) treatment or after 6 h of induction. the indicated strains and conditions. Boxplot elements and statistical analyses are as in Fig. 1e. **b**, Gene expression change of genes categorized into low (grey) middle (white) and high (pink) basal expression bins in cells expressing the indicated constructs. *P* values: two-sided unpaired *t*-test (Benjamini–Hochberg adjusted) of the indicated construct compared to the induced empty plasmid condition (empty). This analysis shows that highly expressed genes are down-regulated in cells expressing 3xFP-NLS while they are up-regulated in cells expressing 3xFP-NES. **c**, Heatmap of mean expression changes for subtelomeric genes across individual and combined (all) chromosomes. Asterisks indicate *P* values determined by comparing expression changes of subtelomeric genes to average expression changes of genes on the rest of the same chromosome (two-sided unpaired *t*-test). **d**, Average Rap1-GFP foci intensity per cell after 6 h mock or induction of the indicated construct. *P* values of comparisons between mock and induced conditions within each strain, calculated via a two-sided *t*-test without adjustment. **e**, Top significantly enriched Cellular Component GO terms for up-regulated genes after 3xFP-NLS induction. Dot size and color represent gene count and adjusted *P* value, respectively. **f**, Correlation between transcript abundance and Pol II occupancy changes. Colors indicate RNA-seq significance (as in Fig. 4a); lines show linear regression fits. **g**, Pol II occupancy changes of genes categorized into low (grey) middle (white) and high (pink) basal expression bins in cells expressing the indicated constructs. Statistics as in **b.** Boxplot elements are as in Fig. 4b.

**Extended Data Figure 5.**
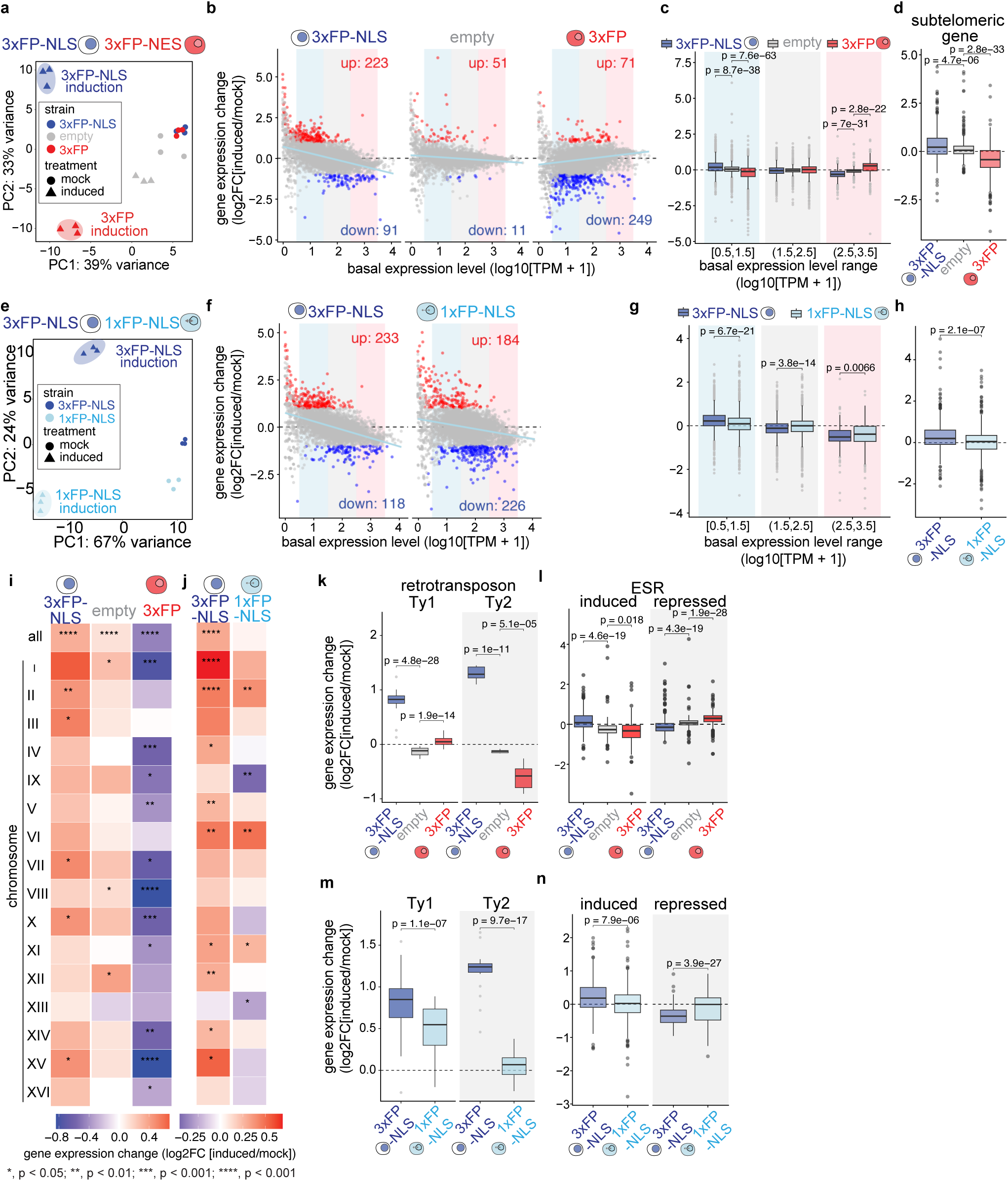
Effect of nuclear transport perturbation on transcriptional reprogramming, related to Figure 4. Panels **a–d**, **i**, **k** and **l** display comparisons between 3xFP-NLS, empty vector, and 3xFP strains. These data show that the simple overexpression of a cytoplasmic enriched protein without a nuclear transport signal is sufficient to cause the same transcriptome changes as expression of the 3xFP-NES construct. Panels **e–h**, **j**, **m** and **n** display comparisons between 3xFP-NLS and 1xFP-NLS strains. These data show that the 3xFP-NLS construct more strongly affects transcription than the 1xFP-NLS construct. Because the 1xFP-NLS imposes a bigger burden on the nuclear import machinery, these data show that the transcriptome changes induced by 3xFP-NLS are not caused by an overload of the nuclear import machinery. **a, e,** PCA analysis of RNA-seq data. **b, f,** Gene expression change (induced vs. mock) plotted against to basal gene expression level (MA plot) determined 6 h after induction of the indicated constructs. Red and blue points indicate significantly up- and down-regulated genes from three replicates (adjusted *P* < 0.05, |log2FC| > 1). Light blue lines show linear regression fits. Shaded backgrounds mark low, medium and high expression bins used for statistical analysis. **c, g,** Comparison of the gene expression changes induced by the different constructs within the indicated low, medium an high expression bins indicated in (**b**) and (**f**). **d, h,** Gene expression changes of subtelomeric genes. **i, j,** Heatmaps representing the gene expression change of subtelomeric genes for individual chromosomes. Asterisks indicate statistical significance in gene expression change between subtelomeric genes and genes in other regions on the same chromosome. **k, m,** Expression changes of Ty1 and Ty2 retrotransposons upon induction of the respective constructs. **l, n,** Expression changes of environmental stress responsive (ESR) genes (stress-induced and stress-repressed) after induction of the respective constructs. Boxplot elements and statistical analyses are as in Fig. 4b.

**Extended Data Figure 6.**
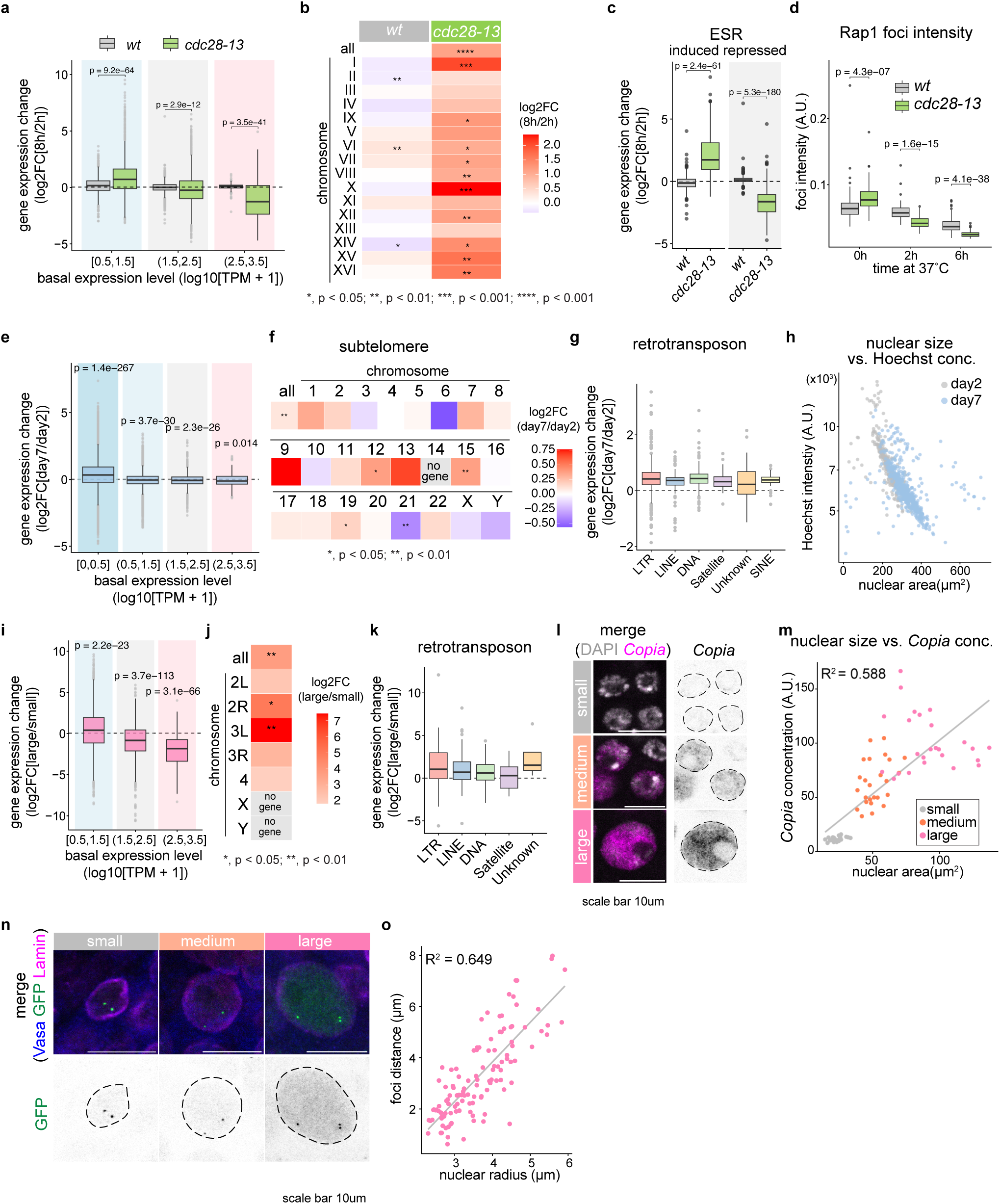
Shared transcriptional reprogramming between excessive cell expansion in G1 arrest and N/C ratio changes, related to Figure 4. **a**-**c**, Re-analysis of gene expression data of *wt* and G1-arrested *cdc28-13* cells grown at 37 °C for 2 h and 8 h from Terhorst et al^48^. **a**, Comparison of the gene expression changes in *wt* and *cdc28-13* cells within the indicated low, medium and high expression bins indicated in Fig. 5b. **b,** Heatmaps representing the gene expression change of subtelomeric genes for individual chromosomes of the indicated strains. Asterisks indicate significance between subtelomeric genes and genes in other regions on the same chromosome. **c,** Expression changes of environmental stress responsive (ESR) genes. **d**, Average Rap1-GFP foci intensity per cell in *wt* and *cdc28-13* grown at 37 °C for the indicated time. *P* values of the comparisons between *wt* and *cdc28-13* strains at each timepoint. **e-g,** Gene expression change in human RPE1-hTERT cells arrested in G1 for 2 days and 7 days using Palbociclib. Expression change across basal expression bins as indicated in Fig. 5h (**e**). Expression change of genes located within 500 kb from chromosome ends on each chromosomes (**f**). *P* values: significance against 0 (one-sample *t*-test). Expression change of different classes of transposable elements (**g**). **h,** Scatter plot showing single-cell nuclear area versus mean Hoechst intensity per nucleus (log10 scale) in human RPE1 cells for the indicated arrest durations. **i**-**k**, Re-analysis of bulk gene expression data comparing differently sized premeiotic cells from *Drosophila* testes (Shi et al)^46^. Expression change across basal expression bins as indicated in Fig. 5n (**i**). Expression change of genes located within 80 kb from chromosome ends on each chromosomes (**j**). *P* values: significance against 0 (one-sample *t*-test). Expression change of different classes of transposable elements (**k**). **l**, Representative high-magnification images of differently sized nuclei in the *Drosophila* testis stained with DAPI and after Fluorescent *In Situ* Hybridization for the RNA of *Copia*, an LTR transposon. Scale bar, 20 µm. **m,** Scatter plot showing single-cell nuclear area versus *Copia* transcript concentration (fluorescent intensity) per cell in *Drosophila* testis. Solid line represents linear regression fit with *R^2^*. **n,** Images of *Drosophila* testis using the LacO/LacI system to the distance between two loci (57A and 60) on chromosome 2. Vasa and Lamin were stained to mark germ cells and nuclear envelopes, respectively. Scale bars, 10µm. Dashed lines: nuclei. **o,** Correlation between nuclear radius and GFP foci distance (*n* = 117) shows that the increasing nuclear size during development is translated into increased inter-chromosome distances. Solid line represents linear regression fit with *R^2^*. Boxplot elements and statistical analyses are as in Fig. 4b.

**Extended Data Figure 7.**
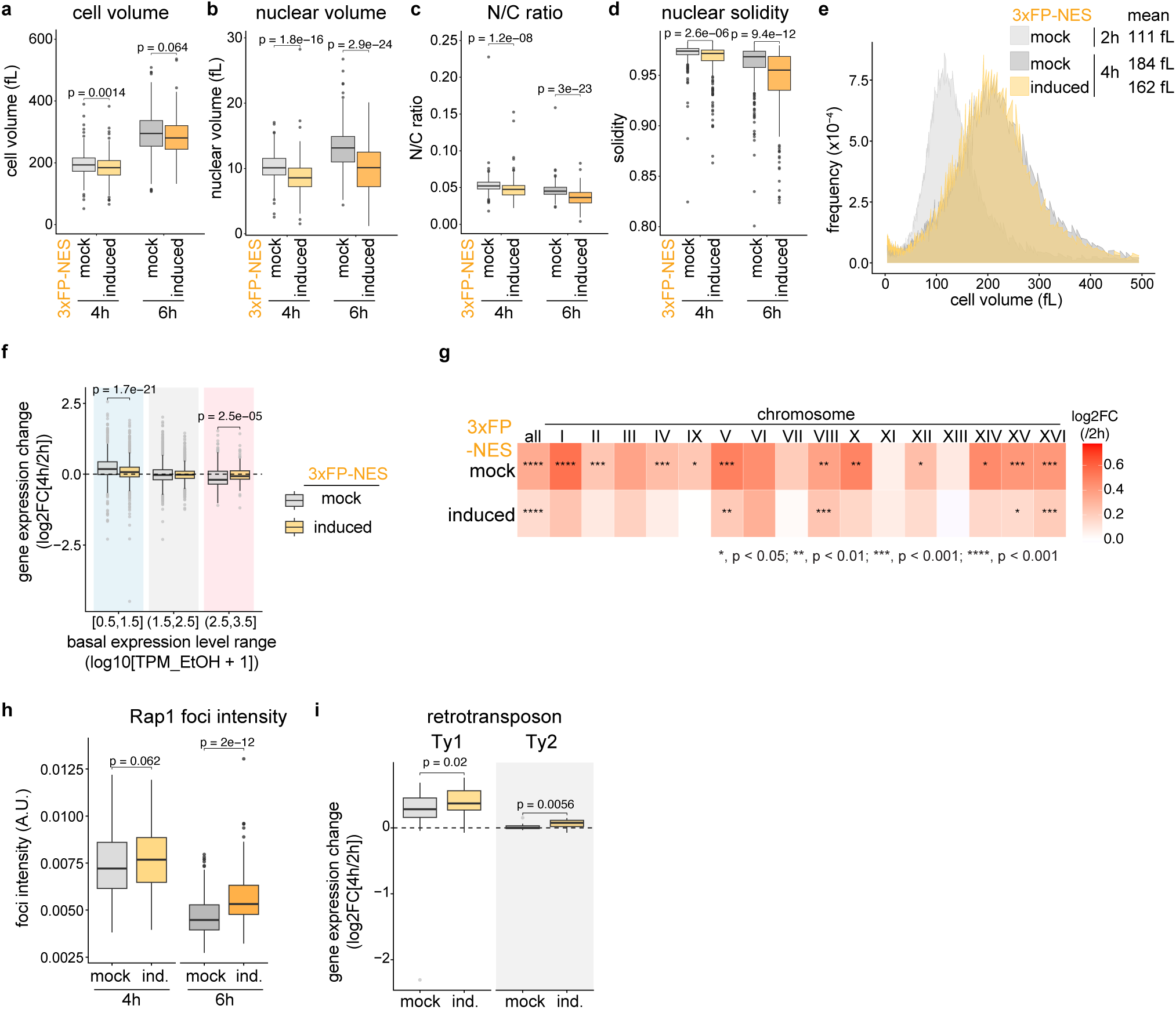
Transcriptome reprogramming in G1-arrest enlarged cells is mitigated by nuclear compression, related to Figure 6. **a–d,** Cell volume (**a**), nuclear volume (**b**), N/C ratio (**c**), and nuclear solidity (**d**) quantified from images of *cdc28-13* cells arrested in G1 at 37 °C with or without induction of 3xFP-NES for 2 hours prior to the shift to 37 °C (4h: mock, *n* = 303, induced, *n* = 349 cells; 6h: mock, *n* = 354, induced, *n* = 247 cells). *P* values via two-sided unpaired *t*-test. **e,** Coulter Counter volume measurements of cell volume distributions over a 4-h time course (2h and 4h) with and without induction of 3xFP-NES. The area plot displays the pooled normalized frequency of cell volumes derived from 3 independent biological replicates per condition. Annotated values indicate the overall mean cell volume (fL) for each condition, calculated as the average of the volume-weighted means across the 3 replicates. **f,** Gene expression changes of G1-arrested *cdc28-13* cells categorized by basal expression bins as indicated in Fig. 6c. Comparison of same arrest conditions with and without induction of 3xFP-NES were performed using an unpaired *t*-test (BH-adjusted) **g,** Heatmap of mean expression changes for subtelomeric genes across individual and combined (all) chromosomes. Asterisks indicate *P* values determined by comparing expression changes of subtelomeric genes to average expression changes of genes on the rest of the same chromosome (two-sided unpaired *t*-test). **h**, Average Rap1-GFP foci intensity per cell in *cdc28-13* cells with and without 3xFP-NES induction after 4 h and 6 h arrest at 37 °C. *P* values indicate comparisons between mock and induced conditions within each strain, calculated via a two-sided *t*-test without adjustment. **i,** Expression change of of Ty1 and Ty2 retrotransposons after 4h G1 arrest. *P* values: paired t-test. Boxplot elements are as in Fig. 4b.

## Supplementary Video Legends

**Supplementary Video 1. Time-lapse imaging of 3xFP-NLS induction.**

Time-lapse fluorescence microscopy of a yeast cell expressing 3xFP-NLS and Nup49-EGFP. The video shows the bright field (BF) and the progressive accumulation of 3xFP-NLS in the nucleus and subsequent transient nuclear envelope rupture events, leading to fluorescent protein leakage into the cytoplasm. Images were acquired every 5 minutes.

## MATERIALS AND METHODS

### Yeast strains and culture

#### Strain information

All *Saccharomyces cerevisiae* strains used in this study are listed in Supplementary Table 1 and are derivatives of the W303 background. Yeast strain construction and genetic modifications were performed using the standard lithium acetate, single-stranded carrier DNA, and PEG transformation protocol^72^. Cells were grown in either YPD medium (1% yeast extract, 2% peptone, 2% glucose) or in synthetic defined (SD) medium (0.67% yeast nitrogen base without amino acids, 2% glucose) supplemented with appropriate amino acids and essential nutrients required for auxotrophic selection as described previously^73^. Unless otherwise stated, all yeast cultures were grown at 30 °C

#### Growth rate and volume measurement

For growth curve analysis, overnight yeast cultures were diluted to OD_600nm_ 0.02 on 48-well plates with the indicated media. Cell growth at 30 °C was monitored by absorbance at 600 nm wavelength every 5 minutes for 48 hours on the plate reader SPECTROstar Nano (BMG Labtech). To measure cell volume, cells from logarithmically growing yeast cells were briefly sonicated, diluted in 10 mL Isotone II buffer (Beckman) and measured on a Coulter Counter (Multisizer 4e, Beckman).

### Human cell lines and culture conditions

HeLa Tet::Cas9 cells (cTT20 cell line) were obtained from the laboratory of Iain Cheeseman^74^. HeLa cells were maintained in Dulbecco’s modified Eagle medium with L-glutamine, 4.5 g/L glucose and sodium pyruvate (ThermoFisher, 41966052), supplemented with 10% heat-inactivated fetal bovine serum and 1% penicillin/streptomycin. Cells were cultured at 37°C with 5% CO_2_. Cell culture conditions and palbociclib treatment of telomerase immortalized human retinal pigment epithelial (RPE1-hTERT) cells were performed as previously described^10^.

#### *Drosophila* strains and culture conditions

All fly strains were raised on standard Bloomington medium and all crosses were maintained at 25°C and 70% humidity. The previously described LacO/LacI system was used to visualize the chromosome 2. A constitutively expressed GFP-tagged bacterial Lac repressor protein binds to two arrays of the Lac operator sequence inserted on the right arm of the second chromosome. As a result, two GFP foci mark two loci on the right arm of the second chromosome. *256xlacO57A 256xlacO60AB/Cyo* (BDSC25375) and *Hsp83-GFP-lacI* (BDSC25376) were obtained from the Bloomington Drosophila Stock Center.

### Nuclear assembly assay in *Xenopus* egg extract

Detailed protocols regarding extract preparation using *Xenopus* egg extracts, nuclear assembly reactions, imaging, and image analysis in this study are described previously^16^. Low-speed/crude (LS) extract and high-speed (HS) extract were used in experiments. The baseline mass concentration (density, mg/mL) of the extracts (surrounding medium) was determined by protein concentration measurement using the Bradford assay. For Low-speed extract preparation, eggs arrested in the metaphase stage were released into interphase by adding CaCl_2_ (0.6 mM), and cyclohexamide (0.1 mg/mL) and energy mix were added to support the assembly reaction. HS extracts were prepared without EGTA/EDTA, allowing endogenous calcium released during egg lysis to drive the extract into interphase. Cycloheximide (0.1 mg/mL) was added prior to the initial crushing spin to arrest the extract in interphase. To the cytosolic layer obtained after ultra-centrifugation (260,000 × *g*) of this extract, salt-washed membranes and energy mix were added. In both assembly systems, Hoechst-33342 (final concentration: 0.05 mg/mL) to visualize DNA was added. Demembranated *Xenopus* sperm nuclei (final concentration: 1000/µL) were subsequently added and incubated at 16–20 °C (with intermediate mixing every 15 min, for 60 min total) to allow nuclear assembly.

### Plasmids

All plasmids used for yeast in this study are listed in Supplementary Tabel 2. Multicopy plasmids are based on the genetic tug-of-war (gTOW) plasmid backbone^30^. A synthetic promoter^29^ as well as different open reading frames were cloned into this plasmid using gene fragment synthesis (Twist Biosciences) and Gibson assembly as described^77^.

### Plasmid accumulation and induction of protein expression

To select and maintain yeast cells harboring many plasmid copies, single colonies were isolated from synthetic defined agar plates lacking uracil (SD-URA) and cultured in liquid SD medium lacking both leucine and uracil (SD-LEU-URA) for a total duration of 48 hours. Throughout this incubation period, cultures were periodically diluted with fresh medium to keep the optical density at 600 nm (OD_600nm_) below 1.0, ensuring continuous exponential growth.

For chemical induction of gene expression, *β*-estradiol (Sigma-Aldrich, E8875) dissolved in ethanol was added to the cultures at a final concentration of 200 nM. For mock-treated control experiments, an equivalent volume of ethanol was added. For G1 synchronization and simultaneous induction of 3xFP-NES in *cdc28-13* temperature-sensitive mutants, *β*-estradiol induction was initiated 2 hours prior to shifting the cultures to the restrictive temperature of 37 °C for G1 arrest.

### Whole protein extraction and quantification

To analyze total cellular protein levels, whole-cell protein extraction was performed using an alkaline pre-treatment method as previously described^78^. Yeast cells equivalent to 2.0 OD_600nm_ units (ca 4x10^7^ cells) were harvested by centrifugation, resuspended in 600 µL 0.1 M NaOH, and incubated at room temperature for 5 minutes. Following centrifugation for 5 minutes at room temperature, the supernatant was discarded. The cell pellet was resuspended in 70 µL of 1× Laemmli SDS sample buffer (62.5 mM Tris-HCl pH 6.8, 2% SDS, 10% glycerol, 5% *β*-mercaptoethanol, 0.01% bromophenol blue) and heat-denatured at 95°C for 5 minutes. The extracted protein samples were diluted 10-fold with 1× Laemmli buffer, and 15 µL (equivalent to approximately 0.043 OD_600_ units of starting cells) was resolved by SDS-PAGE using a Bolt 4–12% Bis-Tris Plus Gel (Thermo Fisher Scientific, NW04120BOX). After electrophoresis, the gel was stained for 30 minutes in Coomassie Brilliant Blue (CBB) staining solution (0.1% CBB R-250 (AppliChem, A3480,0025), 50% methanol, 10% acetic acid) and subsequently destained for 1 hour in destaining solution (40% methanol, 20% acetic acid). Gel images were digitized using an EPSON V700 PHOTO scanner equipped with EPSON Scan software. Densitometric quantification was performed using the "Analyze > Gels" tool in FIJI (ImageJ). The integrated signal intensity of the specific 3xFP band was measured after local background subtraction and normalized against the total integrated intensity of the corresponding entire lane to determine the relative abundance of 3xFP within the whole proteome.

### Microscopy – Sample preparation and image Acquisiion

#### Fluorescent imaging of yeast cells

Cells were plated in 8-well µ-Slide chambers (ibidi, 80821) pre-coated with 2 mg/mL concanavalin A (Con A; Sigma-Aldrich, C2272). Spinning disk confocal imaging of yeast cells was performed on a Nikon Eclipse Ti2 inverted microscope equipped with a Perfect Focus System (PFS) and a Yokogawa CSU-W1 spinning disk confocal scanner unit (10 µm or 50 µm pinhole disk), using a Plan Apo λ 100×/NA 1.45 oil immersion objective. A quad-band dichroic mirror (Di01-T405/488/568/647) was used for laser separation. GFP (or Venus) fluorescence was excited using a 488 nm solid-state laser line and imaged with a 525 nm emission filter (CSU-W1 EM2 Wheel: GFP 525). mCherry fluorescence was excited using a 561 nm laser line and imaged with a 600 nm emission filter (CSU-W1 EM1/EM2 Wheel: Cy3 600). Brightfield images were acquired in widefield transmitted light mode. For each field of view, three-dimensional Z-stack images were collected using a motorized Ti2 Z-drive. Images were captured using a Hamamatsu ORCA-Fusion BT sCMOS camera (C14440-20UP) controlled by the NIS-Elements AR software (Nikon). To induce and maintain G1 arrest in the temperature-sensitive *cdc28-13* mutant, cells were shifted to and imaged at 37 °C.

#### Microfluidic cell culture and Time-lapse imaging

For time-lapse imaging, yeast cells were loaded into a CellASIC ONIX microfluidic plate (EMD Millipore, Y04C-02) according to the manufacturer’s instructions. Cells were continuously perfused with SD-LEU-URA supplemented with either ethanol (used as a vehicle mock control) or *β*-estradiol (final concentration 200 nM; induced) at a constant flow pressure of 10 kPa (∼1.5 psi) at 30 °C throughout the duration of the experiment. Time-lapse widefield fluorescence microscopy was performed using a Nikon Eclipse Ti2 inverted microscope equipped with a Perfect Focus System (PFS) and a Plan Apo λ 60×/NA 1.40 oil immersion objective. Illumination was provided by a Lumencor Spectra/Aura II light engine. GFP fluorescence was excited using the 470 nm line (10% power). mCherry fluorescence was excited using the 575 nm line (10% power). Brightfield images were acquired with a 10 ms exposure (2×2 binning). Excitation and emission were separated using a dual-band dichroic mirror (GFP/mCherry) combined with a Sutter Lambda 10-3 motorized emission filter wheel. For each time point, three-dimensional Z-stack images were collected using a motorized Ti2 Z-drive. Images were captured using a Hamamatsu ORCA-Fusion BT sCMOS camera controlled by the NIS-Elements AR software (version 5.42.03, Nikon). Time-lapse images were acquired every 5 minutes for a total duration of 12 hours.

#### Refractive index tomography in yeast

Samples were imaged using the Tomocube HT-2H quantitative phase imaging (QPI) system equipped with fluorescence imaging capabilities to acquire optical diffraction tomograms at 60X magnification. To correlate the refractive index (RI) with specific structures, fluorescence imaging was performed simultaneously with the RI acquisition. Venus-NLS was imaged using a 470 nm LED and 3xFP was imaged using a 570 nm LED. 2D fluorescence images were acquired in the center plane of the 3D refractive index holotomogram to precisely match the spatial coordinates of the fluorescence signals with the RI map.

#### Optogenetic induction of nuclear export and nuclear size measurement in HeLa cells

HeLa cells stably expressing an NLS–mScarlet3–LEXY probe^55^ were seeded on μ-Slide 4 Well chambers (ibidi, 80426). Prior to imaging, nuclei were stained with SPY650-DNA (Spirochrome, SC501) at a 1:1000 dilution in 1× PBS for 1 h in the dark. Fluorescence images were acquired using a ZEISS Axio Observer 7 epifluorescence microscope equipped with an Axiocam 820 mono camera and controlled by ZEN 3.10 (blue edition). To induce nuclear export of the LEXY probe, cells were exposed to 475-nm blue light for 55 s during each 1-min imaging interval. mScarlet3 and SPY650-DNA fluorescence images were acquired at 1-min intervals for 30 min. Nuclei were identified and segmented based on SPY650-DNA fluorescence.

#### Imaging of nuclear assembly X. laevis egg extract

Imaging of *in vitro* assembled nuclei was performed using a custom-built optical diffraction tomography (ODT) setup based on Mach-Zehnder interferometry and a confocal scanner (Rescan Confocal Microscope, RCM1) installed on the same microscope stand. In ODT, 3D refractive index (RI) tomograms were generated by illuminating the sample at 150 different incident angles using a 532 nm laser and acquiring holograms of scattered light. Simultaneously, by using the same high numerical aperture objective lens (100×, NA 1.3) and the confocal system, correlative fluorescence images of DNA via Hoechst staining were acquired for the same field of view and cell.

#### RPE1 DNA staining and imaging

Cells were washed with 1x PBS prior to 10 min 4% formaldehyde solution fixation. Following fixation, nuclei were stained with Hoechst (0.2 μg/mL Hoechst 33342 in 1x PBS), incubated for 5 min for two times,and wash two time with 1x PBS. Microscopy was performed on a ZEISS Celldiscoverer 7 automated inverted epifluorescence (widefield) microscope using a Plan-Apochromat 20x/NA 0.95 air objective. Hoechst 33258 fluorescence was excited with an integrated 385 nm LED module (370–400 nm excitation) using a 412–433 nm emission filter.

#### Drosophila tissue staining and imaging

Testes from 5 to 7 0-3 day old males were dissected in 1x PBS and transferred to 4% EM-grade paraformaldehyde in PBS and incubated on a nutator for 30 minutes. The fixed tissues were washed three times for 20 minutes each in 1x PBS-T (PBS containing 0.1% Triton-X 100). The tissue was then incubated in 3% Bovine Serum Albumin in 1x PBS-T for 60 minutes for blocking. Primary antibodies were added to the samples and incubated at 4°C overnight. The next day, samples were again washed three times for 15 minutes each with 1x PBS-T and incubated overnight at 4 °C with secondary antibodies diluted in 3% BSA in 1x PBS-T. Samples were again washed as described above on the following day and mounted in VECTASHIELD with DAPI. Following antibodies were used in this study - Rat anti-Vasa (1:100, AB_760351, DSHB), Mouse anti-Lamin Dm0 (1:200; ADL84.12, DSHB). All microscopy images were obtained as described above.

RNA *in situ* hybridization in testes was performed as follows. Testes from 5 to 7 0-3 day old males were dissected in 1x PBS and treated with 4% formaldehyde prepared in RNase free 1x PBS. Post fixation, tissue was washed with 1x PBS two times for 5 mins each, transferred to 70% ethanol diluted in RNase free water and kept on a nutator at 4°C overnight. The following day, the tissue was washed with RNA FISH wash buffer (2xSSC, 10% formamide) for 5 minutes on a nutator at room temperature. The samples were then incubated in hybridization buffer (50 nM probes, 2XSSC, 10% dextran sulfate, 1 g/L *S.cerevisiae* tRNA, 2 mM Vanadyl ribonucleoside complex, 0.5% RNase-free Ultrapure BSA, and 10% deionized formamide) at 37 °C for 12-16 hours. After hybridization, the tissue was washed with RNA FISH wash buffer 2 times at 37 °C for 30 mins each and then mounted in VECTASHIELD with DAPI (Vector Labs). Stellaris *Copia* probes were designed against the LTR consensus sequence using the Stellaris probe designer (LGC Biosearch Technologies). Samples were imaged using a Leica SP8 confocal microscope with a 63x oil-immersion objective (NA=1.4).

### Microscopy analysis

#### Image segmentation and morphological quantification for yeast

All quantitative image analyses, including cell size, nuclear size, and morphological solidity measurements, were performed on maximum intensity projection (Z-max) images using the open-source software Cell-ACDC^79^. Where applicable, fluorescence bleedthrough into the GFP channel caused by high-level 3xFP overexpression was corrected prior to segmentation to ensure boundary accuracy (see the section *"Bleedthrough Correction and Image Demixing"* for detailed methodologies). Two-dimensional (2D) whole-cell segmentation was performed on bright-field Z-max images using the deep learning-based YeaZ v2 model^80^ integrated within Cell-ACDC, followed by manual curation and minor boundary adjustments where necessary. For nuclear segmentation, unbudded cells of interest were manually selected, and their nuclei were segmented on the Venus-NLS fluorescence Z-max channel using an automated Otsu thresholding algorithm^81^. Because segmentation was performed in 2D, three-dimensional (3D) whole-cell and nuclear volumes were estimated from the respective 2D segmentation masks using the longitudinal axis rotational symmetry model ("symmetry rotation") implemented in Cell-ACDC. Furthermore, cell and nuclear solidity (defined as the ratio of the contour area to its convex hull area) were calculated automatically using the standard built-in morphological quantification pipeline in Cell-ACDC.

#### Bleed through correction and image demixing

The high expression level of the red fluorescent proteins caused a significant level of bleed through signal into the GFP channel that was used to segment nuclei and detect Rap1-GFP. To eliminate bleed through contamination from the independent red channel (mCherry) into the dependent green channel (GFP), we implemented a custom pixel-by-pixel demixing pipeline in Python. Cell segmentation masks were first generated from brightfield or phase-contrast images using the YeaZ v2 convolutional neural network. For each image stack, a plane-by-plane background subtraction was performed by calculating the 10th percentile intensity value of each Z-slice and subtracting it from the respective plane. To model the linear bleed through relationship without bias from low-intensity background noise, voxel intensities from the red and green channels within the YeaZ v2 segmentation masks were pooled across reference images. Voxels in the red channel with intensities below a threshold fraction of 0.1 (10% of the maximum intensity) were excluded. A linear regression model (*Y_GFP_* = slope × *X_mC_*_ℎ*erry*_ + intercept) was fitted to the filtered data. To ensure robust signal demixing without over-subtraction, an upward offset corresponding to one residual standard deviation of the fit was applied to the intercept (intercept*_offset_*= intercept + 1.0 × SD*_residual_*). Finally, the predicted bleedthrough contribution was subtracted from the raw green channel pixel-by-pixel;

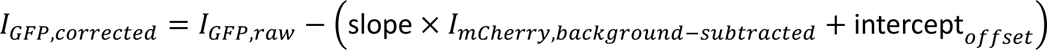

The corrected multi-channel image stacks were exported as OME-TIFF files. All Python scripts and configuration files used for this demixing pipeline are publicly available on GitHub (see *Data and code availability* section).

#### Tomocube analysis for yeast

Cellular and vacuolar regions were segmented using refractive index tomograms. Vacuoles were identified and segmented by exploiting their significantly lower refractive index (RI) compared to the surrounding cytoplasm. Nuclear segmentation was performed based on Venus-NLS fluorescence images. The 3D volumes of the cells and vacuoles were estimated from their 2D segmented regions by approximating their overall shapes as ellipsoids. Specifically, the major radius (*a*) and minor radius (*b*) of each 2D ROI were measured. The volume (*V*) was then calculated using the formula 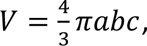, where the unmeasured z-depth radius (*c*) was assumed to be the average of the major and minor radii 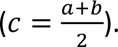 To determine the mass density, small representative ROIs were manually selected within the nucleus and the cytosol using Fiji. The mean intensity of these regions, which corresponds to the absolute RI value, was directly measured. The obtained RI values were subsequently converted to dry-mass density utilizing an RI increment (α) of 0.1907^82^.

#### Nuclear segmentation for nuclear assembly assay in X. laevis egg

When determining nuclear volume, the 3D RI tomograms were used, The nuclei were segmented using either Otsu thresholding or the 3D volume manager, and the volume was calculated by integrating voxels within the segmented region.

#### Image Analysis and Vacuole/Cytosol Segmentation in cdc28-13

To segment the cytosol and exclude vacuolar volumes, we re-analyzed Pgk1-mCherry fluorescence images^2^. Initial pixel classification was performed using ilastik^83^, where pixels were trained and categorized into three classes: cytosol (Pgk1-mCherry), vacuole interior/background, and cell-cell boundaries. The generated multi-scale segmentation outputs were processed using a custom Fiji macro to extract and binarize only the cytosolic regions. These binary masks were then imported into Cell-ACDC, where individual cell IDs were assigned using 3D Otsu thresholding (scaling factor 1.0), followed by manual quality control and corrections. To determine the total cell volume, maximum intensity projection images were segmented using 2D Otsu thresholding in Cell-ACDC. Internal holes corresponding to vacuoles were manually filled via the GUI to create solid cell masks, and cells touching the image edges were excluded from the analysis. Finally, cytosolic volumes were calculated directly from the 3D segmentations, whereas total cell volumes were estimated from the 2D solid masks assuming rotational symmetry in Cell-ACDC. These paired measurements allowed us to determine the relationship between cell size and the vacuole-excluded cytosolic volume (**Extended Data Fig. 3d**). This relationship was then used to derive the theoretical nuclear volume assuming isometric scaling with the cytosol (**Fig. 2h**, blue dashed line).

#### Telomere cluster quantification

For the measurement of telomere clusters labeled by Rap1-GFP, 3D detection and quantification of fluorescent foci (puncta) in Z-stack images were performed using spotMAX software^84^. Taking advantage of the diffuse nucleoplasmic background signal of Rap1-GFP, manually curated 2D nuclear segmentation masks were generated and supplied as spatial boundaries to restrict puncta detection strictly within individual nuclei. Prior to spot identification, input images underwent pre-processing via a Gaussian filter (*σ* = 0.75) and signal sharpening. Fluorescent foci were subsequently detected using a local maximum algorithm (peak_local_max) combined with automated Li thresholding (threshold_li). To enhance spatial precision, spot detection was processed with a 2-fold resolution multiplier in the XY plane. To exclude technical noise, only candidate foci exhibiting a minimum radial dimension of 0.436 µm (6.7 pixels) in XY, an axial span of 1.0 µm (4.0 pixels) in Z, and a segmentation mask of at least 5 voxels were classified as true puncta. For intensity quantification, local background correction was applied using a 5-pixel outer ring surrounding each detected punctum. For every validated focus, we quantified the peak center fluorescence intensity (spot_center_raw_intensity), while overall nuclear volumes (fL) were estimated directly from the 2D nuclear segmentation masks.

#### Image analysis in human cells

For HeLa cells, nuclear size was quantified as the two-dimensional projected area of each segmented nucleus. *For RPE1 cells, n*uclei were segmented based on the Hoechst fluorescence Z-max channel using an automated Otsu thresholding algorithm^81^ and mean intensities were measured with Cell-ACDC^79^.

#### Image analysis for Drosophila testis

All image analysis was performed in FIJI. Z-stacks of the apical tip of the testes were obtained with 0.5 µm slice thickness. Nuclear area and cumulative fluorescence intensity measurements of *Copia* RNA, and DAPI intensity were all obtained by demarcating a Region of Interest (ROI) around individual small, medium and large nuclei. For DAPI intensity, individual pixel values within an ROI were obtained using the pixel histogram function. To reduce variability in fluorescence intensity arising from distance from the objective, we selected cells either in the same slice, or not more than 3 slices apart. For measuring the distance between two GFP foci, cells were selected which had both the foci in the same plane. Nuclear area was measured from the same set of selected cells, and radius values were extracted.

### RNA isolation from yeast and human

#### RNA extraction from yeast cell

Total RNA of yeast cells was extracted using a hot TES-phenol method, followed by column-based cleanup. Briefly, yeast cell pellets containing a defined number of cells were mixed with *Candida albicans* pellets containing a constant number of cells (used as a spike-in control for normalization) were resuspended in TES buffer (10 mM Tris pH 7.5, 10 mM EDTA pH 8.0, 0.5% SDS) and lysed by incubation with saturated phenol at 65°C under continuous agitation. Following centrifugation, the aqueous phase was sequentially re-extracted with saturated phenol and chloroform to remove residual proteins. Total RNA was precipitated from the aqueous phase using 3 M sodium acetate (pH 5.2) and cold isopropanol at -20°C, recovered by centrifugation, washed with 70% ethanol, and resuspended in RNase-free water. To eliminate genomic DNA contamination, the resuspended RNA was treated with DNase I in Buffer RDD (Qiagen, 79254) at room temperature. Following digestion, the RNA was purified, concentrated, and eluted in RNase-free water using the RNeasy MinElute Cleanup Kit (Qiagen, 74204) according to the manufacturer’s instructions. RNA concentration and integrity were assessed using a Qubit Fluorometer (Thermo Fisher Scientific).

#### RNA Extraction from Human RPE1 Cells

Total RNA was extracted from palbociclib-treated RPE1 cells at day 2 (using 1.5 × 10⁶ cells) and day 7 (using 0.5 × 10⁶ cells) of treatment using the RNeasy Mini Kit (Qiagen, 74104) according to the manufacturer’s instructions. Briefly, cell pellets were lysed in 400 µL of Buffer RLT, thoroughly vortexed, and homogenized by centrifugation through a QIAshredder spin column (Qiagen, 79656). The homogenized lysate was combined with ethanol and loaded onto a RNeasy spin column. To eliminate genomic DNA contamination, on-column DNase I digestion was performed using the RNase-Free DNase Set (Qiagen, 79254) in accordance with the manufacturer’s protocol. Following sequential column washes, purified total RNA was eluted in RNase-free water. RNA quantification and quality assessment were performed using a Qubit Fluorometer.

### mRNA sequencing and analysis

mRNA was purified from total RNA using poly-T oligo-attached magnetic beads. Following fragmentation, first-strand cDNA was synthesized using random hexamer primers, followed by second-strand cDNA synthesis. The final cDNA library was completed through end repair, A-tailing, adapter ligation, size selection, amplification, and purification. Library concentration was quantified using a Qubit fluorometer and real-time PCR, while size distribution was assessed using a Bioanalyzer. Quantified libraries were pooled according to their effective concentration and target data amount, and paired-end sequencing NovaSeq X Plus Series (PE150) was performed on an Illumina high-throughput sequencing platform (Novogene Co., Ltd.).

Raw paired-end sequencing reads were processed using fastp (version 0.23.4) for automated adapter trimming and quality filtering (parameters: -q 30 --length_required 20 --detect_adapter_for_pe). Cleaned reads were aligned to a custom combined reference genome—consisting of the *Saccharomyces cerevisiae*reference genome (R64-1-1), the *Candida albicans* reference genome (SC5314, assembly A22), and a custom plasmid sequence encoding 3xFP—using STAR ^85^ in basic two-pass mode (version 2.7.11b; parameters: --twopassMode Basic --outFilterMismatchNmax 3 --chimSegmentMin 2 --alignIntronMax 299999 --outSAMstrandField intronMotif --outSAMtype BAM SortedByCoordinate).

Gene-level read quantification was performed using featureCounts from the package (version 2.16.1)^86^. To accurately quantify transcripts from dense yeast genomes and repetitive regions, multi-mapping reads (up to 10 alignments per read as output by STAR) and reads overlapping multiple genomic features were explicitly retained and fractionally assigned (parameters: isPairedEnd = TRUE, countReadPairs = TRUE, countMultiMappingReads = TRUE, fraction = TRUE, allowMultiOverlap = TRUE). External feature annotations were provided by a custom GTF file combining standard genome annotations with the exogenous sequences. Differential gene expression analysis was conducted using the DESeq2 R package (version 1.52.0)^87^. To account for global shifts in transcription—which would be mathematically masked by standard normalization that assumes constant total RNA levels—we normalized the sequencing depth using the exogenous *C. albicans* spike-in transcripts. Specifically, sample-specific scaling multipliers (size factors) were estimated exclusively using the spike-in transcripts as reference control genes (using the estimateSizeFactors function with the controlGenes argument). These calculated size factors were then applied to normalize the entire endogenous dataset.

#### Re-analysis of yeast *cdc28-13* RNA-seq data

To investigate transcriptome changes during cell size enlargement, we re-analyzed previous RNA-seq dataset^48^. Specifically, single-end sequencing reads corresponding to wild-type (WT) and temperature-sensitive *cdc28-13* mutant strains exposed to 37°C for 2 hours and 8 hours (3 biological replicates per condition, 12 samples in total; NCBI SRA accessions SRR22926215–SRR22926293) were retrieved and fasterq-dump from the SRA Toolkit (version 3.0.3). The bioinformatic pipeline followed the same general workflow as described above, with modifications tailored for single-end reads and standard reference annotations. Specifically, automated quality filtering and adapter trimming were performed using fastp without paired-end detection flags. The cleaned single-end reads were aligned exclusively to the standard *Saccharomyces cerevisiae* reference genome (R64-1-1) using STAR in basic two-pass mode, and read counting was executed with featureCounts set to single-end mode (isPairedEnd = FALSE), while retaining the multi-mapping rescue and fractional assignment parameters (countMultiMappingReads = TRUE, fraction = TRUE, allowMultiOverlap = TRUE) to ensure consistent quantification of repetitive regions. To eliminate technical artifacts and library size distortions common in public datasets, ribosomal RNAs, tRNAs, small nuclear RNAs, and small nucleolar RNAs (specifically genes matching the prefixes *RDN, YLR154, YLR162, YLR157, YLR159, YLR156, ETS, ITS, snR, SCR1,* and *LSR1*) were strictly filtered out prior to downstream analyses.

### Transcriptomic and Transposable Element (TE) Analysis in Human Cells

For the RNA-seq analysis of human retinal pigment epithelial (RPE1) cells, raw paired-end sequencing reads were trimmed and quality-filtered using fastp (requiring -q 30 and --length_required 20). Cleaned reads were aligned to the human reference genome (GRCh38/hg38) using STAR in basic two-pass mode. To accurately capture transcription from highly repetitive sequences and transposable elements (TEs), the alignment parameters were modified to permit up to 100 multi-mapping locations per read (--outFilterMultimapNmax 100, --winAnchorMultimapNmax 100).

To quantify both protein-coding genes and repetitive elements simultaneously, aligned BAM files were first name-sorted using multi-threaded samtools (version 1.19.2; samtools sort -n), followed by quantification using TEcount from the TEtranscripts software package (version 2.2.4; in multi-mode, --mode multi)^88^. Annotation files consisted of the GENCODE primary assembly (v44) for standard genes and the corresponding RepeatMasker GTF file (GRCh38_GENCODE_rmsk_TE.gtf) for transposable elements.

For differential expression analysis using DESeq2, raw count matrices were filtered to retain genes and TEs that exhibited at least 1 count in a minimum of 4 independent samples (rowSums(counts >= 1) >= 4).

#### Re-analysis of *Drosophila* RNA-seq data

To investigate transcriptomic and TE dynamics across different cell sizes in *Drosophila melanogaster*, we re-analyzed publicly available single-end RNA-seq data from public dataset^50^ (NCBI SRA accession number SRP239059). Specifically, single-end sequencing reads corresponding to isolated small cells (replicates S4 and S8) and large cells (replicates LC1 and LC2) were downloaded using prefetch and fasterq-dump. The bioinformatic pipeline followed the exact same architecture as described for our human dataset, with parameters optimized for single-end reads and *Drosophila* genome annotations. Quality filtering and adapter trimming were conducted using fastp in single-end mode (-q 30, --length_required 20). Cleaned reads were mapped to the *D. melanogaster* reference genome (BDGP6.32/dm6) using STAR in basic two-pass mode, permitting up to 100 multi-mapping locations per read (--outFilterMultimapNmax 100, --winAnchorMultimapNmax 100) to capture repetitive TE transcription. BAM files were pre-sorted by read name using multi-threaded samtools (samtools sort -n) and subsequently quantified using TEcount (TEtranscripts package in multi-mode, --mode multi) against the Ensembl release 109 GTF for standard genes and the BDGP6 RepeatMasker GTF (BDGP6_rmsk_TE.gtf) for transposable elements. For differential expression analysis comparing large versus small cells using DESeq2, count matrices were filtered to retain features with at least 1 count in a minimum of 3 samples (rowSums(counts > 1) >= 3).

### ChIP sequencing

#### Chromatin immunoprecipitation (ChIP)

Prior to cross-linking, *Saccharomyces cerevisiae* cells were mixed with *Candida glabrata* cells at a 1:2 ratio by cell number to serve as a spike-in control for normalization. Cells were cross-linked with 1% formaldehyde for 15 minutes at 30°C and quenched with 0.125 M glycine. Harvested cell pellets (250–350 mg) were resuspended in FA lysis buffer (50 mM HEPES-KOH pH 8.0, 150 mM NaCl, 1 mM EDTA pH 8.0, 1% Triton X-100, 0.1% sodium deoxycholate) freshly supplemented with 1 mM PMSF and protease inhibitor cocktail (Thermo Fisher Scientific, 78429), and disrupted using ceramic beads in a FastPrep Bead Beating Systems (MP Biomedicals). After centrifugation (13,500 rpm, 10 minutes, 4°C), chromatin was sheared by bioruptor (diagenode) to yield fragment sizes between 100-300 bp, and clarified by centrifugation (15,000 rpm, 5 minutes, 4°C).

Sheared chromatin was incubated for 5 hours to overnight at 4°C with Protein G Dynabeads (Thermo Fisher Scientific, 10003D) that were blocked with 0.5% BSA, after rinsed with PBS, were pre-coated with Anti-RNA Polymerase II Antibody (anti Rpb1, Merck, clone 8WG16) antibody, and then rinsed with PBS once. After incubation with the sheared chromatin, beads were sequentially washed twice with FA lysis buffer, twice with high salt FA lysis buffer (50 mM HEPES-KOH pH 8.0, 500 mM NaCl, 1 mM EDTA pH 8.0, 1% Triton X-100, 0.1% sodium deoxycholate), twice with ChIP wash buffer (10 mM Tris-HCl pH 8.0, 1 mM EDTA, 0.25 M LiCl, 0.5% NP-40, 0.5% sodium deoxycholate), and rinsed once with TE wash buffer (10 mM Tris-HCl pH 8.0, 1 mM EDTA, 50 mM NaCl). Chromatin was eluted in 1% SDS elution buffer at 65°C for 15 minutes. Cross-links of both ChIP and input samples were reversed at 65°C overnight, followed by sequential digestion with RNase A (Merck, 10109169001; 37°C, 1 hour) and Proteinase K (AppliChem, A3830,0100; 65°C, 2 hours). DNA was purified using the ChIP DNA Clean & Concentrator Kit (Zymo Research, D5201) and quantified using a Qubit Fluorometer.

#### Library preparation and sequencing

Prior to library preparation, the quality and concentration of ChIP and input DNA samples were assessed using a TapeStation 4200 system with D1000 ScreenTapes and analyzed via TapeStation Analysis Software v5.2 (Agilent Technologies). Sequencing libraries were prepared from 4.0 ng of input or immunoprecipitated DNA using the NEBNext Ultra II DNA Library Prep Kit for Illumina (New England Biolabs, E7645), strictly following the manufacturer’s protocol. Briefly, following end-prep and adapter ligation, the reactions were purified using a 0.9× volume of SPRI beads without size selection. Adapter-ligated DNA fragments were subsequently enriched by PCR and subjected to a final post-amplification cleanup using a 0.9× bead ratio. Final library concentrations and fragment size distributions were validated on the TapeStation 4200 system. The resulting libraries were equimolar-pooled and sequenced on an Illumina NovaSeq X Plus platform (Illumina) using a high-output 500M flow cell in a 2 × 150 bp paired-end configuration, according to the manufacturer’s standard clustering and sequencing protocols

#### Chip-seq data analysis

Raw paired-end sequencing reads were processed using fastp (version 0.23.4) for automated adapter trimming and quality filtering (parameters: -q 30, --length_required 20, --detect_adapter_for_pe). Cleaned paired-end reads were aligned to a custom combined reference genome—consisting of the *Saccharomyces cerevisiae* reference genome (R64-1-1), the *Candida glabrata* reference genome (CBS138), and a custom plasmid sequence (3xFP)— using bowtie2 (version 2.5.2)^89^. The resulting alignments were converted, sorted by coordinate, and indexed using samtools. To ensure robust data interpretation, a single outlier replicate (3xFP-NES mock replicate 2) exhibiting technical variance was systematically excluded from all downstream analyses and visualization.

Signal Normalization and Differential Occupancy Analysis Genome-wide signal coverage files (BigWig format) were generated using the deeptools (version 3.5.6) suite. Normalized ChIP enrichment was first computed as Counts Per Million (CPM) using bamCoverage (parameters: --normalizeUsing CPM, --centerReads, --binSize 10). To evaluate treatment-induced changes in occupancy, log2 fold change (log2FC) BigWig files comparing immunoprecipitated (IP) samples against their corresponding input controls were generated using bamCompare (--scaleFactorsMethod readCount, --operation log2, --binSize 10). For gene-level quantification, a custom annotation BED file (Sc_gene.bed) was constructed by standard Ensembl GTF gene coordinates (0-based). Gene-body enrichment scores were extracted from the log_2_ FC BigWig files using multiBigwigSummary BED-file. Statistical analyses were performed in R using dplyr and rstatix. For each gene, differential occupancy between estradiol (induced) and ethanol (mock) treatments was evaluated using two-sided Welch’s two-sample *t*-tests (paired = FALSE, var.equal = FALSE), followed by Benjamini-Hochberg false discovery rate (FDR) adjustment (padj). To account for global signal shifts between strains and conditions, raw log_2_ FC values were normalized by subtracting the median log_2_FC across all genes for each strain (median-centering).

Metagene Profiling and Genomic Track Visualization Metagene profiles across transcription start sites (TSS) and transcription end sites (TES) were generated using computeMatrix scale-regions in deeptools (parameters: -b 500 -a 500 --regionBodyLength 1000 --binSize 10). Within R, to visualize occupancy dynamics across different transcription levels, genes were ranked by their basal Pol II occupancy in the EtOH mock condition and stratified into three percentiles: Top 10%, Mid 10–50%, and Bottom 50%. Each replicate matrix was median-centered across all genomic bins prior to calculating pooled bin averages. The absolute Pol II occupancy ratio for each sample was calculated by normalizing the target read counts (*S. cerevisiae*genome + plasmid) to the spike-in (*C. glabrata*) read counts, and subsequently normalizing the IP ratio against the corresponding Input ratio. The calculation was performed using the following formula:

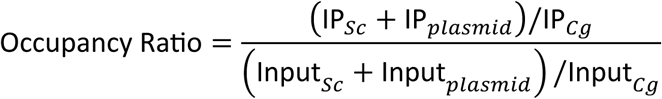

### Statistics and reproducibility

#### Sample sizes (n) and experimental replication

No statistical methods were used to pre-determine sample sizes; rather, sample sizes were chosen based on standard experimental practices in the field to ensure adequate statistical power and reproducibility. Where indicated, n represents the number of individual cells/nuclei examined or the number of independent biological replicates performed. Experimental replication structures across all figures are categorized as follows:

##### Biological replicates from distinct pre-cultures

For global transcriptomic profiling, genomic occupancy (ChIP-seq), and growth/protein assays, data were derived from 3 independent biological replicates started from separate pre-cultures (Fig. 4a, b, e–m; Fig. 6b–e; Extended Data Fig. 1a, b; Extended Data Fig. 2f; Extended Data Fig. 4a–c, e-g; Extended Data Fig. 5a–n; Extended Data Fig. 7f, g, i). The mRNA-seq data of human RPE1 cells was derived from 4 independent biological replicates started from separate pre-culture (Fig. 5g–j; Extended Data Fig. 6e–g).

##### Pooled datasets from independent experiments

For quantitative single-cell imaging analyses requiring large-scale modeling or volume-dependent stratification, data were pooled from 3 or 4 independent biological experiments (Fig. 2c, e-h; Fig. 3d; Fig. 5l, r; Extended Data Fig. 3a; Extended Data Fig. 6h, m, o; Extended Data Fig. 7e). Data for *X. laevis in vitro* nuclear assembly assays were similarly pooled from multiple extract preparations (Fig. 1o, p).

##### Representative experiments

To evaluate single-cell distribution dynamics and nuclear morphometrics without batch-to-batch visual artifacts, data are shown for 1 representative experiment out of 2 or 3 independent biological experiments performed, with the remaining independent experiments exhibiting identical quantitative and phenotypic trends (Fig. 1c, d, e–h, j, k, n; Fig. 3c; Fig. 4c, d; Fig. 5e, f, k, q; Fig. 6f, g; Extended Data Fig. 1c, e–i; Extended Data Fig. 2a, b, d, e, g–l; Extended Data Fig. 4d; Extended Data Fig. 6d, l, n; Extended Data Fig. 7a–d, h).

##### Re-analysis of public datasets

For comparisons involving excessive cell expansion, public RNA-seq and imaging datasets were re-analyzed following our standardized bioinformatics pipeline. This includes data from *cdc28-13* yeast (Fig. 5b–d; Extended Data Fig. 6a–c) and *Drosophila* spermatogenesis (Fig. 5n–p; Extended Data Fig. 6i–k).

##### Exact cell numbers

The exact number of analyzed cells/nuclei (*n*) for time-lapse imaging, volume quantifications, and in vitro nuclear assembly assays are detailed directly in the respective figure legends (e.g., Fig. 1e-h, k, o, p; Fig. 2c, e, f, h; Fig. 3d; Fig. 4d; Fig. 5f, l, r; Fig. 6g; Extended Data Fig. 2g-l).

#### Statistical analyses and hypothesis testing

Statistical analyses were performed using R and SciPy. All statistical tests applied were two-sided unless otherwise specified.

##### Parametric comparisons

Comparisons between two experimental groups were evaluated using two-sided unpaired Student’s t-tests (Fig. 1e–h, o, p; Fig. 3d; Fig. 4.l; Extended Data Fig. 3a; Extended Data Fig. 4b–d, g; Extended Data Fig. 5c, g, i, j; Extended Data Fig. 6a, b, d; Extended Data Fig. 7a–d, f–h). Where appropriate to compare paired experimental features or matched reference controls within specific datasets, two-sided paired t-tests were utilized (Fig. 4b, e, f, h–j; Fig. 5c, d; Fig. 6d, e; Extended Data Fig. 5d, h, k–n; Extended Data Fig. 6c; Extended Data Fig. 7i). For single-group comparisons against a hypothesized null value, one-sample t-tests were applied against a reference value of 1.0 (Fig. 4m) or 0 (Fig. 5i, o; Extended Data Fig. 6e, f, i, j).

##### High-throughput sequencing analyses

Statistical significance for differential gene expression was determined using two-sided Wald tests. *P* values were adjusted for multiple hypothesis testing using the Benjamini–Hochberg false discovery rate (FDR/BH) method, defining adjusted P < 0.05 and |log2FC| > 1 as statistically significant (Fig. 4a, g; Fig. 5b, h, j, n, p; Fig. 6c; Extended Data Fig. 5b, f).

##### Regression and correlation

Linear regression models were fitted to evaluate relationships between continuous variables, with model fits displayed as solid or light blue lines (Fig. 4a, g; Fig. 5b, h, n; Fig. 6c; Extended Data Fig. 4f; Extended Data Fig. 5b, f; Extended Data Fig. 6m, o). Smoothed conditional mean regressions were applied for volume and morphometric scaling correlations (Fig. 2h; Extended Data Fig. 3d).

#### Data presentation and visualization

In all boxplots presented (Fig. 1e–h; Fig. 2f; Fig. 3d; Fig. 4b, e, f, h–j; Fig. 5c, d, i, o; Fig. 6d, e; Extended Data Fig. 2g-l; Extended Data Fig. 3a; Extended Data Fig. 4b, d, g; Extended Data Fig. 5c, d, g, h, k–n; Extended Data Fig. 6a, c, d, e, g, i, k; Extended Data Fig. 7a-d, f, h, i), the center line represents the median, box boundaries indicate the 25th and 75th percentiles (interquartile range, IQR), and whiskers extend up to 1.5× the IQR, with individual data points beyond the whiskers plotted as outliers. Continuous time-course measurements (nuclear size in HeLa cells and *X. laevis* cell), mRNA concentrations, and global Pol II occupancy are presented as the mean ± s.e.m. (standard error of the mean) across replicates or cells (Fig. 1k, o, p; Fig. 4l, m; Extended Data Fig. 1i). Bar plot for protein abundance proportion and continuous time-course growth curves are presented as the mean ± s.d. (standard deviation) across biological replicates (Extended Data Fig. 1b; Extended Data Fig. 2f). Principal component analysis (PCA) was performed using VST-normalized counts to assess sample clustering and overall data variance (Fig. 6b; Extended Data Fig. 4a; Extended Data Fig. 5a, e).

#### Data normalization and exclusion

##### Normalization

mRNA-seq counts were normalized via variance-stabilizing transformation (VST; Fig. 6b; Extended Data Fig. 4a; Extended Data Fig. 5a, e) or expressed as standard transcripts per million (TPM) / counts per million (CPM) (Fig. 4a, b, e, f; Fig. 5b-d, h-j, n-p; Fig. 6c–e; Extended Data Fig. 4b, c, e, f, g; Extended Data Fig. 5b–d, f–h, k–n; Extended Data Fig. 6a-c, e-g, i-k; Extended Data Fig. 7f, g, i). To explicitly account for global transcriptional shifts and estimate total cellular mRNA concentrations, read counts were normalized using *C. albicans* spike-in controls (Fig. 4l). For ChIP-seq, raw read counts were first normalized to sequencing depth as counts per million (CPM). To evaluate relative occupancy and distribution profiles, these normalized signals were subsequently median-centered where appropriate (Fig. 4g–k). In contrast, global Pol II occupancy was absolute-quantified utilizing *C. glabrata* spike-in reads (Fig. 4m).

##### Data exclusion

In accordance with established quality-control standards, two specific sequencing samples exhibiting extreme technical variance and acting as significant statistical outliers on principal component analysis (PCA) were systematically excluded prior to downstream differential quantification: (1) In the genome-wide Pol II ChIP-seq dataset (Fig. 4g–k, m), one replicate (3xFP-NES mock replicate 2) was excluded from the 3 biological replicates initiated from separate pre-cultures. (2) In the nuclear compression time-course RNA-seq dataset (Fig. 6b–e; Extended Data Fig. 7f–i), one replicate (4h induced replicate 2) was excluded from the 3 biological replicates initiated from separate pre-cultures. No cells, imaging data points, or other biological replicates were excluded unless failing automated segmentation quality controls.

#### Reproducibility and randomization

All independent biological replicates, representative imaging findings, and quantitative phase imaging (QPI) profiles yielded reproducible results across repeated trials. Randomization and blinding were not applicable to this study, as experimental conditions (e.g., specific genetic modifications, light illumination, and chemical inductions) required explicit identification for appropriate sample execution, grouping, and bioinformatics pipeline processing.

## SUPPLEMENTARY TABLES

**Supplementary Table 1.** List of *S. cerevisiae* strains used in this study.

| strain name | genotype | description | Reference |
| --- | --- | --- | --- |
| GN3225 | *W303, MATa, ade2-1, leu2-3, ura3, trp1-1, his3-11,15, can1-100, GAL, psi+, Nup49-GFP:KAN, LexA-ER-B112:HIS3, [LexA(4)-mScarlet-mCherry2x-NLS-pTOW, URA3, leu2-89]* | 3xFP-NLS<br>Nup49-GFP | This study |
| GN3229 | *W303, MATa, ade2-1, leu2-3, ura3, trp1-1, his3-11,15, can1-100, GAL, psi+, Nup49-GFP:KAN, LexA-ER-B112:HIS3, [pTOW40836, URA3, leu2-89]* | empty<br>Nup49-GFP | This study |
| GN3506 | *W303, MATa, ade2-1, leu2-3, ura3, trp1-1, his3-11,15, can1-100, GAL, psi+, Nup49-GFP:KAN, LexA-ER-B112:HIS3 [LexA(4)-mScarlet-mCherry2x-NES-pTOW, URA3, leu2-89]* | 3xFP-NES<br>Nup49-GFP | This study |
| GN448 | *W303, MATa, ade2-1, leu2-3, ura3, trp1-1, his3-11,15, can1-100, GAL, psi+, Nup49-GFP:KAN, LexA-ER-B112:HIS3 [LexA(4)-mScarlet-mCherry2x-pTOW, URA, leu2-89]* | 3xFP Nup49-GFP | This study |
| GN1684 | *W303, MATa, ade2-1, leu2-3, ura3, trp1-1, his3-11,15, can1-100, GAL, psi+, Nup49-GFP:KAN, LexA-ER-B112:HIS3, [LexA(4)-mCherry-NLS-pTOW, URA3, leu2-89]* | 1xFP Nup49-GFP | This study |
| GN3537 | *W303, MATa ade2-1, leu2-3, ura3, trp1-1, his3-11,15, can1-100, GAL, psi+, LexA-ER-B112:HIS3, pCTS1-2xVenus-SV40NLS:URA3, [LexA(4)-mScarlet-2xmCherry-NLS-pTOW, URA3, leu2-89]* | 3xFP-NLS<br>Venus-NLS | This study |
| GN3548 | *W303, MATa ade2-1, leu2-3, ura3, trp1-1, his3-11,15, can1-100, GAL, psi+, LexA-ER-B112:HIS3, pCTS1-2xVenus-SV40NLS:URA3, [LexA(4)-mScarlet-mCherry2x-NES-pTOW, URA3, leu2-89]* | 3xFP-NES<br>Venus-NLS | This study |
| GN3516 | *W303, MATa, ade2-1, leu2-3, ura3, trp1-1, his3-11,15, can1-100, GAL, psi+ Nup49-mCherry:NAT LexA-ER-B112:HIS3 GFP-RAP:ADE2* | Rap1-GFP | This study |
| GN3585 | *W303, MATa, ade2-1, leu2-3, ura3, trp1-1, his3-11,15, can1-100, GAL, psi+ Nup49-mCherry:NAT LexA-ER-B112:HIS3 GFP-RAP:ADE2, [LexA(4)-mScarlet-mCherry2x-NLS-pTOW, URA3, leu2-89]* | Rap1-GFP<br>3xFP-NLS | This study |
| GN3586 | *W303, MATa, ade2-1, leu2-3, ura3, trp1-1, his3-11,15, can1-100, GAL, psi+ Nup49-mCherry:NAT LexA-ER-B112:HIS3 GFP-RAP:ADE2, [LexA(4)-mScarlet-mCherry2x-NES-pTOW, URA3, leu2-89]* | Rap1-GFP<br>3xFP-NES | This study |
| GN3581 | *W303, MATa, ade2-1, leu2-3, ura3, trp1-1, his3-11,15, can1-100, GAL, psi+ Nup49-mCherry:NAT LexA-ER-B112:HIS3 GFP-RAP:ADE2 cdc28-13* | Rap1-GFP<br>cdc28-13 | This study |
| GN3728 | *W303, MATa, ade2-1, leu2-3, ura3, trp1-1, his3-11,15, can1-100, GAL, psi+ Nup49-mCherry:NAT LexA-ER-B112:HIS3 GFP-RAP:ADE2, [LexA(4)-mScarlet-mCherry2x-NES-pTOW, URA3, leu2-89] cdc28-13* | Rap1-GFP<br>cdc28-13<br>3xFP-NES | This study |
| GN3654 | *W303, MATa ade2-1, leu2-3, ura3, trp1-1, his3-11,15, can1-100, GAL, psi+, LexA-ER-B112:HIS3, pCTS1-2xVenus-SV40NLS:URA3, [LexA(4)-mScarlet-mCherry2x-NES-pTOW, URA3, leu2-89] cdc28-13* | cdc28-13<br>3xFP-NES<br>Venus-NLS | This study |

**Supplementary Table 2.** List of plasmids used in this study.

| strain name | genotype | description | Reference |
| --- | --- | --- | --- |
| pGN376 | PACT1(-1-520)-LexA-ER-haB112-TCYC1 | LEXA system | Ottoz et al<sup>29</sup> |
| pGN440 | URA3, leu2-89/2μ | empty | Moriya et al<sup>90</sup> |
| pGN470 | LexA(4)-mScarlet-2xmCherry-NLS, URA3, leu2-89/2μ | 3xFP-NLS | This study |
| pGN548 | LexA(4)-mScarlet-2xmCherry-NES, URA3, leu2-89/2μ | 3xFP-NES | This study |
| pGN477 | LexA(4)-mScarlet-2xmCherry,URA3, leu2-89/2μ | 3xFP | This study |
| pGN504 | LexA(4)-mCherry,URA3, leu2-89/2μ | 1xFP-NLS | This study |

## SUPPLEMENTARY NOTES

### Supplementary Note 1: Basic mathematical model for estimating the total number of colloidal particles (***N_t_***)

We constructed a mathematical model to estimate the total number of colloid osmotically active particles in the cell (*N_t_*) based on the experimentally obtained N/C ratios (*r*) and vacuole volume fractions (*φ_v_*). We are using here some extensive variables (number, volume) for ease of interpretability, but the model can be rescaled to intensive variables (concentrations, volume ratios). In particular, the extensive version allows us to naturally express results as the behavior of an “average” cell. Let *r*_0_ and *φ_v_*_,0_ be the initial N/C ratio and vacuolar fraction prior to 3xFP induction, respectively, and *r_i_* and *φ_v_*_,*i*_ (*i* for *induced*) be the corresponding observed values after the induction of *n* 3xFP molecules. Based on Van’t Hoff’s law, Young-Laplace’s law, and hydrodynamic simulations^21,91^, the relationship between the number of colloidal particles in the nucleus and cytoplasm (*N_nuc_* and *N_cyto_*), their respective volumes (*V_nuc_* and *V_cyto_*), the nuclear radius (*R^nuc^*), the nuclear membrane tension (*σ*), and the Boltzmann constant (*k_B_*) is described as:

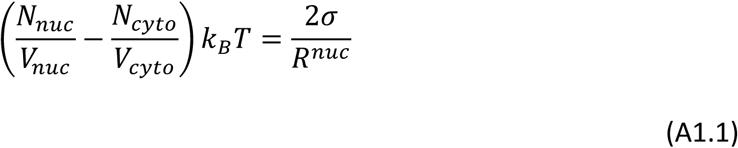

Assuming negligible nuclear membrane tension (*σ* = 0), and substituting *N_cyto_* = *N_t_* − *N_nuc_* and *V_cyto_* = *V_cell_* − *V_nuc_* − *V_vac_* into this equation, we rearrange it for the N/C ratio (*V_nuc_*/*V_cell_*) to obtain:

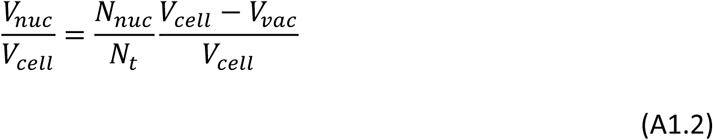

In the case where V_vac_ = 0,

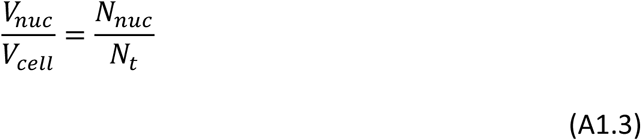

Next, without loss of generality (WLOG), we set the total cell volume *V_cell_* = 1 to shift from absolute volumes (*V*) to experimentally measurable volume fractions (*r* and *φ_v_*). We also define the fraction of nuclear proteins as *R_n_* = *N_nuc_*/*N_t_*. Consequently, in the hypothetical absence of a vacuole (*φ_v_* = 0), *R_n_* simply equals the N/C ratio. This relationship is given by:

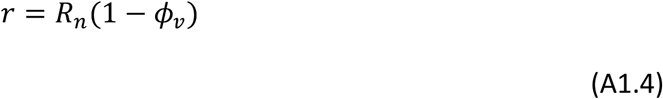

When considering protein induction, we assume that the number fraction partitioning of the endogenous proteome would remain unchanged – that is, if e.g. 8% of the particles were localizing to the nucleus before induction, 8% of the endogenous particles would localize to the nucleus after induction.

The change in the nuclear protein distribution ratio before and after 3xFP induction (Δ*R_n_*) is defined as Δ*R_n_* = *R_n_*_,*i*_ − *R_n_*_,0_. When *n* new 3xFP particles are induced, let *f* denote the fraction of these particles targeted to the nucleus (*f* = 1 for complete nuclear localization, and *f* = 0 for complete cytoplasmic localization). If we fix *N_t_*, the induced proteins are *n*/*N_t_*, and the remaining (*N_t_* − *n*)/*N_t_* are partitioned as before, the effective change in the number of nuclear colloidal particles (Δ*N_nuc_*) is defined as the number of particles forcibly translocated to the nucleus (*n* ⋅ *f*) minus the number of particles that would have naturally partitioned into the nucleus according to the initial distribution ratio (*n* ⋅ *R_n_*_,0_):

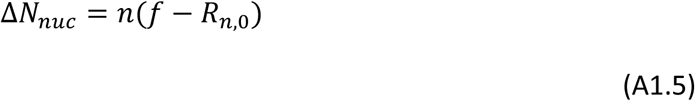

Since the change in the distribution ratio (Δ*R_n_*) is equal to the fraction of effectively added particles (Δ*N_nuc_*/*N_t_*), we have Δ*R_n_* = Δ*N_nuc_*/*N_t_*. Substituting the aforementioned equations and solving for *N_t_* yields:

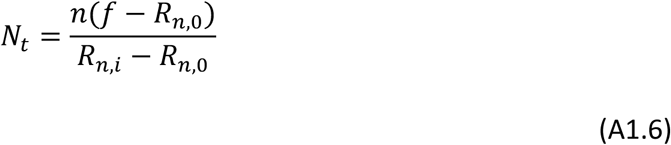

Using the relationship 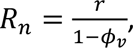, we derive the general equation for *N_t_* consisting solely of measurable experimental parameters:

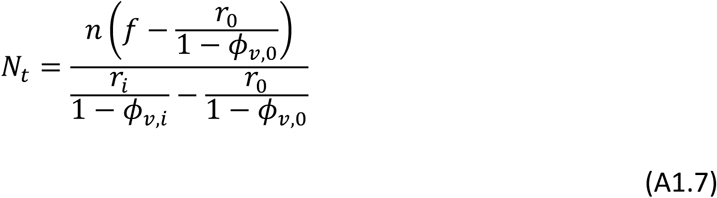

By solving this general equation for the final N/C ratio (*r_i_*), the observed N/C ratio after 3xFP-NLS and 3xFP-NES induction can be expressed as a function of the total number of colloid particles (*N_t_*), the number of induced 3xFP molecules (*n*), the vacuole fractions before and after induction (*φ_v_*_,0_ and *φ_v_*_,*i*_), and the initial N/C ratio (*r*_0_). For the 3xFP-NLS system (*f* = 1), the equation rearranges to:

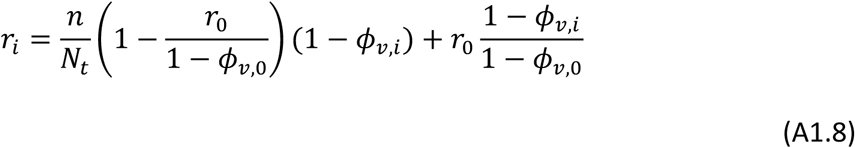

Similarly, for the 3xFP-NES system (*f* = 0), the equation simplifies to: (2)

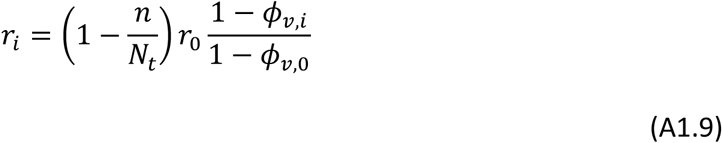

Note that this works well because by using the experimentally available 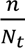 (a function of the proteome mass fraction that is induced) and the induced and uninduced ratios, we can recover *N_t_* for a cell of arbitrary size.

### Supplementary Note 2: Quantification of absolute 3xFP molecule numbers from fluorescence intensity

To convert the relative fluorescence intensity of 3xFP measured at the single-cell level into absolute molecule numbers, first, we determined the mass fraction of 3xFP (*f_p_*) relative to the total proteome for both 3xFP-NLS and 3xFP-NES expressing strains by CBB staining. Because CBB staining intensity is generally proportional to protein mass, the mass fraction was quantified as the ratio of the specific 3xFP band signal to the total signal of the entire lane. Assuming a total cellular protein mass (*M*) of 5 pg (BNID 106225)^92^, the average absolute copy number of 3xFP per cell (*n_av_*_g_) was calculated using the Avogadro constant (*N_A_*) and the known molecular weight of 3xFP (*MW_p_* = 8.8 × 10^4^ g/mol) according to the following equation:

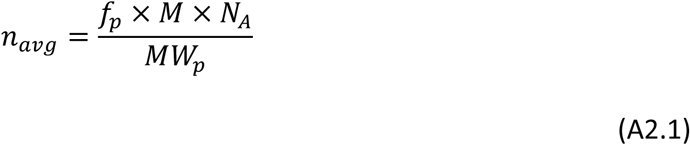

Parallel to the biochemical quantification, single-cell fluorescence imaging was performed on identically induced cell populations. For each cell, the mean fluorescence intensity was extracted, and the local auto-background signal was subtracted. To robustly correlate the fluorescence signal with the biochemical data, we calculated a volume-weighted average fluorescence intensity (*I_av_*_g_) across the induced 3xFP-NLS cell population by integrating the total fluorescence intensity and total cellular volume across all biological replicates. A unicersal conversion factor (*α*) representing the absolute number of molecules per fluorescence intensity unit was then derived by dividing the biochemically calculated average copy number by this volume-weighted average intensity (*α* = *n_av_*_g_/*I_av_*_g_). Finally, the absolute number of 3xFP molecules in each individual cell (*n_i_*) —for both the 3xFP-NLS and 3xFP-NES expressing strains—was computationally determined by multiplying its specific background-subtracted fluorescence intensity (*I_i_*) by the corresponding construct-specific conversion factor:

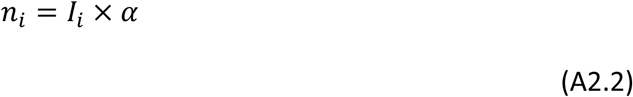

Detailed descriptions of the image processing pipelines, along with the raw datasets and the complete R scripts used for these conversions, are provided in the GitHub (https://github.com/shinohsawa/Ohsawa_et_al_2026).

### Supplementary Note 3: Estimation of the effective number of colloidal particles based on protein abundance

To estimate the effective number of osmotically active colloidal particles in the cell, we progressively filtered the proteome dataset and corrected for protein complex formation. First, we utilized the whole-organism protein abundance dataset from PaxDb[https://pax-db.org]^33^ for Saccharomyces cerevisiae alongside protein mass data. Assuming a total cellular protein mass of 5 pg (BNID 106225)^92^, we calculated the absolute copy number of each protein using its abundance (in ppm) and the weighted average molecular weight derived from the entire proteome. At this stage, all proteins were initially assumed to exist as independent monomers to determine the theoretical maximum number of particles. Next, because intracellular proteins frequently assemble into multiprotein complexes that act as single colloidal entities, we integrated stoichiometry data from the Complex Portal database^93^. To do this, complex abundances are calculated as the median of the abundances of their subunits (scaled by their stoichiometry in the complex). Proteins with unknown stoichiometry were assigned a stoichiometric index of 1. This was also used to annotate an estimated mass of the complex. Subunits of the complexes are then removed from this version of the proteome. For proteins not described in the Complex Portal, particularly homo-oligomers (e.g., homodimers, trimers, and tetramers), we utilized the “Subunit structure” annotations from the UniProt Knowledgebase (UniProtKB)^94^ to adjust their effective particle counts. Additional specific stoichiometric adjustments were manually applied to cytoskeletal proteins and nucleosome. Specifically, two-thirds of actin was assumed to be polymerized with only the remaining one-third existing as soluble monomers^95^, 90% of tubulin was considered polymerized^96^,and nucleosomes were excluded because they are bound within the immobile chromatin matrix and do not act as freely diffusing colloidal particles^24^. This yielded an effective particle count closely reflecting the physiological state, consisting of both unassociated monomers and assembled complexes. To define macromolecules that meaningfully contribute to the colloidal osmotic pressure in the cyto-nucleoplasm, we then applied a size exclusion threshold, removing small protein particles (both monomers and complexes) with a molecular mass of 40 kDa or less. Finally, to isolate the soluble colloidal particles freely diffusing within the cyto-nucleoplasmic fluid, we utilized Cellular Component annotations from Gene Ontology (GO_C). Proteins and complexes associated with “membrane” or “vacu” (vacuolar components) were excluded from the dataset. Through these progressive refinements and exclusions, we quantified the final effective number of colloidal particles physically acting as independent osmotically active entities within the intracellular space. Furthermore, to analyze the dynamic changes during cell size increase, we utilized time-series proteomics data from a *cdc28-13* mutant^2^. We anchored our baseline absolute particle counts to the 1 h time point and applied the relative fold-changes from the proteomics data to estimate the final effective number of colloidal particles at all subsequent time points. Using Gene Ontology Cellular Component (GO_C) annotations, we then categorized these particles and calculated their relative proportions across four subcellular fractions: cytoplasm, nucleus, both (shared), and other. Detailed descriptions of the data processing procedures, along with the raw datasets and the complete R script used for this analysis, are available in the GitHub (https://github.com/shinohsawa/Ohsawa_et_al_2026).

### Supplementary Note 4: Extended mathematical model for estimating nuclear envelope tension (***σ***)

While Supplementary Note 1 derives the total number of particles (*N_t_*) under the assumption of negligible nuclear envelope tension (*σ* = 0), empirical data revealed a discrepancy in the estimated *N_t_* values between the 3xFP-NLS and 3xFP-NES systems. Furthermore, observations of nuclear envelope rupture indicate that nuclear expansion, particularly in the 3xFP-NLS system, is physically constrained by membrane tension. To resolve this discrepancy, we extend our model by returning to the fundamental equation presented in Supplementary Note 1 and re-evaluating the nuclear envelope tension (*σ*) that was previously assumed to be zero. For simplicity, we assume the vacuolar volume fraction is zero (*φ_v_* = 0) both before and after induction, meaning the N/C ratio equals the nuclear volume fraction (*r*_0_ and *r_i_* for before and after induction, respectively). This is justified by the observation, detailed in the main text, that the the experimental values of *φ_v_* would introduce a much smaller correction than the 3xFP induction effect. We also assume that the cell nucleus in its initial steady state possesses sufficient excess membrane area (wrinkles), such that the initial nuclear envelope tension prior to induction is zero (*σ*_0_ = 0). In the induced state, the relationship between the colloid osmotic pressure balance across the nuclear envelope and the membrane tension is described by the fundamental equation from Supplementary Note 1:

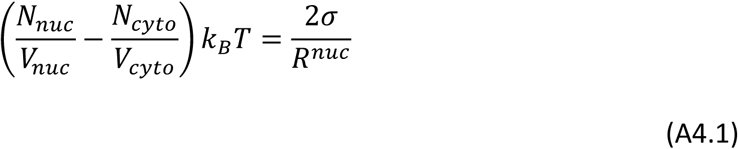

Considering the system upon 3xFP-NLS induction (*f* = 1), the effective change in the number of nuclear particles is defined as Δ*N_nuc_* = *n*(1 − *r*_0_), as established in Supplementary Note 1. Consequently, the number of particles in the nucleus (*N_nuc_*) and the cytoplasm (*N_cyto_*), based on total particle conservation (*N_cyto_* = *N_t_* − *N_nuc_*), can be expressed as *N_nuc_* = *N_t_r*_0_ + *n*(1 − *r*_0_) and *N_cyto_* = *N_t_*(1 − *r*_0_) − *n*(1 − *r*_0_). Let the absolute total cell volume be *V_cell_*, making the respective compartmental volumes *V_nuc_* = *V_cell_r_i_* and *V_cyto_* = *V_cell_*(1 − *r_i_*). Substituting these particle numbers and compartmental volumes into the fundamental equation, rearranging the concentration difference on the left side over a common denominator, and subsequently expressing *V_cell_r_i_* back as *V_nuc_* yields the following relationship:

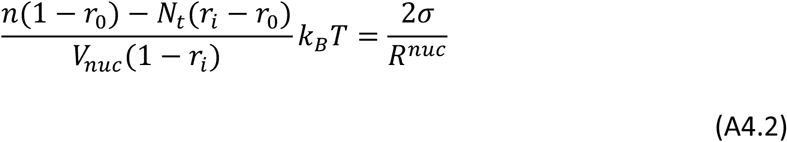

To solve this equation for *σ*, we express the nuclear radius directly in terms of the nuclear volume (*V_nuc_*) using the geometric formula for the volume of a sphere, 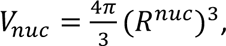, which gives:

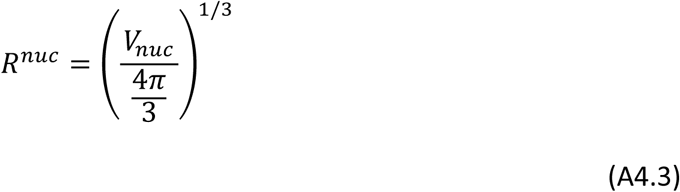

By substituting this geometric relationship and rearranging the terms, we derive the general equation to estimate the nuclear envelope tension solely from measurable experimental parameters:

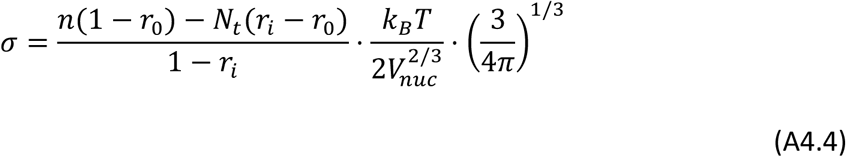

To practically calculate the nuclear envelope tension (*σ*) in the 3xFP-NLS system using this equation, the true total number of particles (*N_t_*) is required as a baseline reference. In the 3xFP-NES system, osmotically active particles accumulate in the cytoplasm, driving water efflux from the nucleus and causing it to shrink. During this shrinking and relaxation process, no physical expansion tension is generated on the nuclear envelope; it merely develops wrinkles or slack. Therefore, the assumption that the tension is zero (*σ* = 0) can be applied to the 3xFP-NES system. Consequently, the *N_t_* value estimated from the 3xFP-NES system based on the model in Supplementary Note 1 can be considered the true total number of colloidal particles in the cell. By substituting this 3xFP-NES-derived *N_t_* into the equation above, we can quantitatively derive the tension *σ* exerted on the nuclear envelope due to the forced particle translocation in the 3xFP-NLS system.

## Notes

### Competing Interest Statement

The authors have declared no competing interest.

## REFERENCE

1. Brandt, A. et al. Developmental Control of Nuclear Size and Shape by *kugelkern* and *kurzkern*. Curr. Biol. 16, 543–552 (2006).

2. Neurohr, G. E. et al. Excessive Cell Growth Causes Cytoplasm Dilution And Contributes to Senescence. Cell 176, 1083–1097.e18 (2019).

3. Newport, J. & Kirschner, M. A major developmental transition in early *Xenopus* embryos: I. characterization and timing of cellular changes at the midblastula stage. Cell 30, 675–686 (1982).

4. Newport, J. & Kirschner, M. A major developmental transition in early *Xenopus* embryos: II. control of the onset of transcription. Cell 30, 687–696 (1982).

5. Prabha, H., Phan, P. & Levy, D. L. Getting nuclear size just right – emerging mechanisms regulating nuclear scaling and morphology. J. Cell Sci. 139, jcs264825 (2026).

6. Boveri, T. Zellenstudien V. Über die Abhängigkeit der Kerngrösse und Zellenzahl bei Seeigellarven von der Chromosomenzahl der Ausganszellen. Jenaische Z. Für Naturwissenschaft 39, 445–524 (1905).

7. Hertwig, R. Ueber die Korrelation von Zell-und Kerngrösse und ihre Bedeutung für die Geschlechtliche Differenzierung und die Teilung der Zelle. Biol. Cent. 23, 49–62 (1903).

8. Bateman, J. R. & Anderson, D. J. Taming the giant within. PLOS Genet. 15, e1008098 (2019).

9. Edens, L. J., White, K. H., Jevtic, P., Li, X. & Levy, D. L. Nuclear size regulation: from single cells to development and disease. Trends Cell Biol. 23, 151–159 (2013).

10. Manohar, S. et al. Genome homeostasis defects drive enlarged cells into senescence. Mol. Cell 83, 4032–4046.e6 (2023).

11. Jorgensen, P. et al. The Size of the Nucleus Increases as Yeast Cells Grow. Mol. Biol. Cell 18, 3523–3532 (2007).

12. Neumann, F. R. & Nurse, P. Nuclear size control in fission yeast. J. Cell Biol. 179, 593–600 (2007).

13. Willis, L. et al. Cell size and growth regulation in the *Arabidopsis thaliana* apical stem cell niche. Proc. Natl. Acad. Sci. 113, (2016).

14. Estrada, M. E. et al. Non-coding DNA dictates cell size and impairs fitness by sequestering RNA polymerase. Preprint at 10.64898/2026.07.10.737727 (2026).

15. Shuter, B. J., Thomas, J. E., Taylor, W. D. & Zimmerman, A. M. Phenotypic Correlates of Genomic DNA Content in Unicellular Eukaryotes and Other Cells. Am. Nat. 122, 26–44 (1983).

16. Biswas, A. et al. Conserved nucleocytoplasmic density homeostasis drives cellular organization across eukaryotes. Nat. Commun. 16, 7597 (2025).

17. Maeshima, K., Iino, H., Hihara, S. & Imamoto, N. Nuclear size, nuclear pore number and cell cycle. Nucleus 2, 113–118 (2011).

18. Cantwell, H. & Nurse, P. Unravelling nuclear size control. Curr. Genet. 65, 1281–1285 (2019).

19. Jevtić, P., Edens, L. J., Vuković, L. D. & Levy, D. L. Sizing and shaping the nucleus: mechanisms and significance. Curr. Opin. Cell Biol. 28, 16–27 (2014).

20. Kume, K. et al. A systematic genomic screen implicates nucleocytoplasmic transport and membrane growth in nuclear size control. PLOS Genet. 13, e1006767 (2017).

21. Lemière, J., Real-Calderon, P., Holt, L. J., Fai, T. G. & Chang, F. Control of nuclear size by osmotic forces in *Schizosaccharomyces pombe*. eLife 11, e76075 (2022).

22. Mukherjee, R. N., Chen, P. & Levy, D. L. Recent advances in understanding nuclear size and shape. Nucleus 7, 167–186 (2016).

23. Levy, D. L. & Heald, R. Mechanisms of intracellular scaling. Annu. Rev. Cell Dev. Biol. 28, 113–135 (2012).

24. Deviri, D. & Safran, S. A. Balance of osmotic pressures determines the nuclear-to-cytoplasmic volume ratio of the cell. Proc. Natl. Acad. Sci. 119, e2118301119 (2022).

25. Finan, J. D. & Guilak, F. The effects of osmotic stress on the structure and function of the cell nucleus. J. Cell. Biochem. 109, 460–467 (2010).

26. Rollin, R., Joanny, J.-F. & Sens, P. Physical basis of the cell size scaling laws. eLife 12, e82490 (2023).

27. Mohr, D., Frey, S., Fischer, T., Güttler, T. & Görlich, D. Characterisation of the passive permeability barrier of nuclear pore complexes. EMBO J. 28, 2541–2553 (2009).

28. Mitchison, T. J. Colloid osmotic parameterization and measurement of subcellular crowding. Mol. Biol. Cell 30, 173–180 (2019).

29. Ottoz, D. S. M., Rudolf, F. & Stelling, J. Inducible, tightly regulated and growth condition-independent transcription factor in *Saccharomyces cerevisiae*. Nucleic Acids Res. 42, e130–e130 (2014).

30. Moriya, H., Makanae, K., Watanabe, K., Chino, A. & Shimizu-Yoshida, Y. Robustness analysis of cellular systems using the genetic tug-of-war method. Mol. Biosyst. 8, 2513–2522 (2012).

31. Kafri, M., Metzl-Raz, E., Jona, G. & Barkai, N. The Cost of Protein Production. Cell Rep. 14, 22–31 (2016).

32. Janssen, A. F. J., Breusegem, S. Y. & Larrieu, D. Current Methods and Pipelines for Image-Based Quantitation of Nuclear Shape and Nuclear Envelope Abnormalities. Cells 11, 347 (2022).

33. Huang, Q., Szklarczyk, D., Oehninger, J. & von Mering, C. PaxDb v6.0: reprocessed, LLM-selected, curated protein abundance data across organisms. Nucleic Acids Res. 54, D427–D439 (2026).

34. Lemière, J., Ren, Y. & Berro, J. Rapid adaptation of endocytosis, exocytosis, and eisosomes after an acute increase in membrane tension in yeast cells. eLife 10, e62084 (2021).

35. Rawicz, W., Olbrich, K. C., McIntosh, T., Needham, D. & Evans, E. Effect of Chain Length and Unsaturation on Elasticity of Lipid Bilayers. Biophys. J. 79, 328–339 (2000).

36. Chen, Y., Huang, J.-H., Phong, C. & Ferrell, J. E. Viscosity-dependent control of protein synthesis and degradation. Nat. Commun. 15, 2149 (2024).

37. Delarue, M. et al. mTORC1 Controls Phase Separation and the Biophysical Properties of the Cytoplasm by Tuning Crowding. Cell 174, 338–349.e20 (2018).

38. Uhler, C. & Shivashankar, G. V. Regulation of genome organization and gene expression by nuclear mechanotransduction. Nat. Rev. Mol. Cell Biol. 18, 717–727 (2017).

39. Gotta, M. et al. The clustering of telomeres and colocalization with Rap1, Sir3, and Sir4 proteins in wild-type *Saccharomyces cerevisiae*. J. Cell Biol. 134, 1349–1363 (1996).

40. Hozé, N., Ruault, M., Amoruso, C., Taddei, A. & Holcman, D. Spatial telomere organization and clustering in yeast *Saccharomyces cerevisiae* nucleus is generated by a random dynamics of aggregation–dissociation. Mol. Biol. Cell 24, 1791–1800 (2013).

41. Kothiwal, D. & Laloraya, S. A SIR-independent role for cohesin in subtelomeric silencing and organization. Proc. Natl. Acad. Sci. 116, 5659–5664 (2019).

42. Ruault, M., De Meyer, A., Loïodice, I. & Taddei, A. Clustering heterochromatin: Sir3 promotes telomere clustering independently of silencing in yeast. J. Cell Biol. 192, 417–431 (2011).

43. Curcio, M. J., Lutz, S. & Lesage, P. The Ty1 LTR-Retrotransposon of Budding Yeast, *Saccharomyces cerevisiae*. Microbiol. Spectr. 3, 3.2.19 (2015).

44. Brauer, M. J. et al. Coordination of Growth Rate, Cell Cycle, Stress Response, and Metabolic Activity in Yeast. Mol. Biol. Cell 19, 352–367 (2008).

45. Gasch, A. P. et al. Genomic Expression Programs in the Response of Yeast Cells to Environmental Changes. Mol. Biol. Cell 11, 4241–4257 (2000).

46. Swaffer, M. P. et al. RNA polymerase II dynamics and mRNA stability feedback scale mRNA amounts with cell size. Cell 186, 5254–5268.e26 (2023).

47. Lanz, M. C. et al. Increasing cell size remodels the proteome and promotes senescence. Mol. Cell 82, 3255–3269.e8 (2022).

48. Terhorst, A. et al. The environmental stress response regulates ribosome content in cell cycle-arrested *S. cerevisiae*. Front. Cell Dev. Biol. 11, 1118766 (2023).

49. Fuller, M. T. Genetic control of cell proliferation and differentiation in *Drosophila* spermatogenesis. Semin. Cell Dev. Biol. 9, 433–444 (1998).

50. Shi, Z. et al. Single-cyst transcriptome analysis of *Drosophila* male germline stem cell lineage. Development 147, dev184259 (2020).

51. Keber, F. C., Nguyen, T., Mariossi, A., Brangwynne, C. P. & Wühr, M. Evidence for widespread cytoplasmic structuring into mesoscale condensates. Nat. Cell Biol. 26, 346–352 (2024).

52. Narduzzi, G. et al. Weak interactions drive selective proteome demixing and tune the differential response to environmental perturbations. Preprint at 10.64898/2026.07.22.739830 (2026).

53. Park, J. O. et al. Metabolite concentrations, fluxes and free energies imply efficient enzyme usage. Nat. Chem. Biol. 12, 482–489 (2016).

54. Cantwell, H. & Nurse, P. A systematic genetic screen identifies essential factors involved in nuclear size control. PLOS Genet. 15, e1007929 (2019).

55. Moriizumi, H. et al. Nuclear size is genetically controlled and influences cell fate. Preprint at 10.64898/2026.08.26.747275 (2026).

56. Lemière, J., Tan, Z. & Chang, F. Nuclear size and physical properties of the nucleoplasm are determined by colloid osmotic pressure at the nuclear envelope. Preprint at 10.64898/2026.07.21.739918 (2026).

57. Chen, Y., Zhao, G., Zahumensky, J., Honey, S. & Futcher, B. Differential Scaling of Gene Expression with Cell Size May Explain Size Control in Budding Yeast. Mol. Cell 78, 359–370.e6 (2020).

58. Miller, K. E., Vargas-Garcia, C., Singh, A. & Moseley, J. B. The fission yeast cell size control system integrates pathways measuring cell surface area, volume, and time. Curr. Biol. 33, 3312–3324.e7 (2023).

59. Schmoller, K. M., Turner, J. J., Kõivomägi, M. & Skotheim, J. M. Dilution of the cell cycle inhibitor Whi5 controls budding-yeast cell size. Nature 526, 268–272 (2015).

60. Lanz, M. C. et al. Genome dilution by cell growth drives starvation-like proteome remodeling in mammalian and yeast cells. Nat. Struct. Mol. Biol. 31, 1859–1871 (2024).

61. De Cecco, M. et al. L1 drives IFN in senescent cells and promotes age-associated inflammation. Nature 566, 73–78 (2019).

62. Gorbunova, V. et al. The role of retrotransposable elements in ageing and age-associated diseases. Nature 596, 43–53 (2021).

63. Li, W. et al. Activation of transposable elements during aging and neuronal decline in Drosophila. Nat. Neurosci. 16, 529–531 (2013).

64. Van Steensel, B., Smogorzewska, A. & De Lange, T. TRF2 Protects Human Telomeres from End-to-End Fusions. Cell 92, 401–413 (1998).

65. Lawlor, M. A., Cao, W. & Ellison, C. E. A transposon expression burst accompanies the activation of Y-chromosome fertility genes during *Drosophila* spermatogenesis. Nat. Commun. 12, 6854 (2021).

66. Bonaccorsi, S., Pisano, C., Puoti, F. & Gatti, M. Y chromosome loops in *Drosophila* melanogaster. Genetics 120, 1015–1034 (1988).

67. Klumpe, S. et al. In-cell structure and snapshots of *copia* retrotransposons in intact tissue by cryo-ET. Cell 188, 2094–2110.e18 (2025).

68. Fingerhut, J. M., Moran, J. V. & Yamashita, Y. M. Satellite DNA-containing gigantic introns in a unique gene expression program during *Drosophila* spermatogenesis. PLOS Genet. 15, e1008028 (2019).

69. Venkei, Z. G. et al. A maternally programmed intergenerational mechanism enables male offspring to make piRNAs from Y-linked precursor RNAs in *Drosophila*. Nat. Cell Biol. 25, 1495–1505 (2023).

70. Ozata, D. M., Gainetdinov, I., Zoch, A., O’Carroll, D. & Zamore, P. D. PIWI-interacting RNAs: small RNAs with big functions. Nat. Rev. Genet. 20, 89–108 (2019).

71. Czech, B. et al. piRNA-Guided Genome Defense: From Biogenesis to Silencing. Annu. Rev. Genet. 52, 131–157 (2018).

72. Gietz, R. D. & Schiestl, R. H. High-efficiency yeast transformation using the LiAc/SS carrier DNA/PEG method. Nat. Protoc. 2, 31–34 (2007).

73. Dunham, M. J., Gartenberg, M. R. & Brown, G. M. Methods in Yeast Genetics and Genomics. (Cold Spring Harbor laboratory press, Cold Spring Harbor (N.Y.), 2015).

74. McKinley, K. L. et al. The CENP-L-N Complex Forms a Critical Node in an Integrated Meshwork of Interactions at the Centromere-Kinetochore Interface. Mol. Cell 60, 886–898 (2015).

75. Robinett, C. C. et al. In vivo localization of DNA sequences and visualization of large-scale chromatin organization using lac operator/repressor recognition. J. Cell Biol. 135, 1685–1700 (1996).

76. Vazquez, J., Belmont, A. S. & Sedat, J. W. The Dynamics of Homologous Chromosome Pairing during Male *Drosophila* Meiosis. Curr. Biol. 12, 1473–1483 (2002).

77. Gibson, D. G. et al. Enzymatic assembly of DNA molecules up to several hundred kilobases. Nat. Methods 6, 343–345 (2009).

78. Kushnirov, V. V. Rapid and reliable protein extraction from yeast. Yeast 16, 857–860 (2000).

79. Padovani, F., Mairhörmann, B., Falter-Braun, P., Lengefeld, J. & Schmoller, K. M. Segmentation, tracking and cell cycle analysis of live-cell imaging data with Cell-ACDC. BMC Biol. 20, 174 (2022).

80. Dietler, N. et al. A convolutional neural network segments yeast microscopy images with high accuracy. Nat. Commun. 11, 5723 (2020).

81. Otsu, N. A Threshold Selection Method from Gray-Level Histograms. IEEE Trans. Syst. Man Cybern. 9, 62–66 (1979).

82. Zhao, H., Brown, P. H. & Schuck, P. On the Distribution of Protein Refractive Index Increments. Biophys. J. 100, 2309–2317 (2011).

83. Berg, S. et al. ilastik: interactive machine learning for (bio)image analysis. Nat. Methods 16, 1226–1232 (2019).

84. Padovani, F. et al. SpotMAX: a generalist framework for multi-dimensional automatic spot detection and quantification. Preprint at 10.1101/2024.10.22.619610 (2024).

85. Dobin, A. et al. STAR: ultrafast universal RNA-seq aligner. Bioinformatics 29, 15–21 (2013).

86. Liao, Y., Smyth, G. K. & Shi, W. featureCounts: an efficient general purpose program for assigning sequence reads to genomic features. Bioinformatics 30, 923–930 (2014).

87. Love, M. I., Huber, W. & Anders, S. Moderated estimation of fold change and dispersion for RNA-seq data with DESeq2. Genome Biol. 15, 550 (2014).

88. Jin, Y., Tam, O. H., Paniagua, E. & Hammell, M. TEtranscripts: a package for including transposable elements in differential expression analysis of RNA-seq datasets. Bioinformatics 31, 3593–3599 (2015).

89. Langmead, B. & Salzberg, S. L. Fast gapped-read alignment with Bowtie 2. Nat. Methods 9, 357–359 (2012).

90. Moriya, H., Shimizu-Yoshida, Y. & Kitano, H. In Vivo Robustness Analysis of Cell Division Cycle Genes in *Saccharomyces cerevisiae*. PLoS Genet. 2, e111 (2006).

91. Wu, C.-H., Fai, T. G., Atzberger, P. J. & Peskin, C. S. Simulation of Osmotic Swelling by the Stochastic Immersed Boundary Method. SIAM J. Sci. Comput. 37, B660–B688 (2015).

92. Von Der Haar, T. & McCarthy, J. E. G. Intracellular translation initiation factor levels in *Saccharomyces cerevisiae* and their role in cap-complex function. Mol. Microbiol. 46, 531–544 (2002).

93. Balu, S. et al. Complex portal 2025: predicted human complexes and enhanced visualisation tools for the comparison of orthologous and paralogous complexes. Nucleic Acids Res. 53, D644–D650 (2025).

94. The UniProt Consortium et al. UniProt: the Universal Protein Knowledgebase in 2025. Nucleic Acids Res. 53, D609–D617 (2025).

95. Gonzalez Rodriguez, S., Wirshing, A. C. E., Goodman, A. L. & Goode, B. L. Cytosolic concentrations of actin binding proteins and the implications for in vivo F-actin turnover. J. Cell Biol. 222, e202306036 (2023).

96. Winey, M. & Bloom, K. Mitotic Spindle Form and Function. Genetics 190, 1197–1224 (2012).

